# S-adenosylmethionine synthases specify distinct H3K4me3 populations and gene expression patterns during heat stress

**DOI:** 10.1101/2022.03.30.486419

**Authors:** Adwait A. Godbole, Sneha Gopalan, Thien-Kim Nguyen, Alexander Munden, Paula Vo, Caroline A. Lewis, Jessica B. Spinelli, Thomas G. Fazzio, Amy K. Walker

## Abstract

Methylation is a widely occurring modification that requires the methyl donor S-adenosylmethionine (SAM) and acts in regulation of gene expression and other processes. SAM is synthesized from methionine, which is imported or generated through the 1-carbon cycle (1CC). Alterations in 1CC function have clear effects on lifespan and stress responses, but the wide distribution of this modification has made identification of specific mechanistic links difficult. Exploiting a dynamic stress-induced transcription model, we find that two SAM synthases in *Caenorhabditis elegans*, SAMS-1 and SAMS-4, contribute differently to modification of H3K4me3, gene expression and survival. We find that *sams-4* enhances H3K4me3 in heat shocked animals lacking *sams-1*, however, *sams-1* cannot compensate for *sams-4*, which is required to survive heat stress. This suggests that the regulatory functions of SAM depend on its enzymatic source and that provisioning of SAM may be an important regulatory step linking 1CC function to phenotypes in aging and stress.

## Introduction

The 1-Carbon cycle (1CC) is a group of interconnected pathways that link essential nutrients such as methionine, folate and vitamin B12 to the production of nucleotides, glutathione, and S-adenosylmethionine (SAM), the major methyl donor ^1^ (**Fig1A**). SAM is important for the production of polyamines and phosphatidylcholine (PC), a methylated phospholipid, and is also essential for the methylation of RNA, DNA and proteins such as histones ^2^. Thus, 1CC connects nutrients with the production of a key cellular regulator of epigenetic function, SAM.

**Figure 1.**
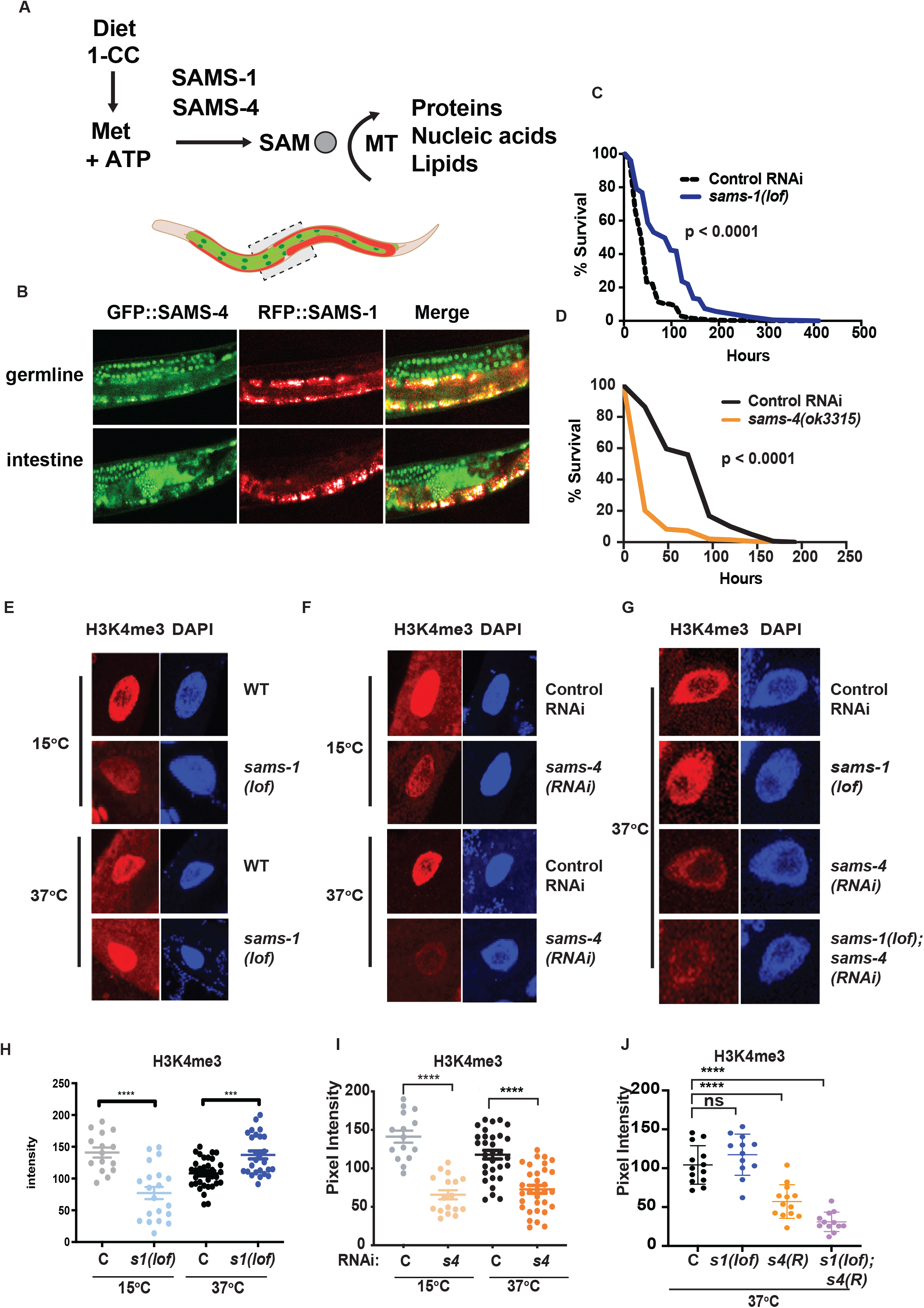
*sams-1*-independent acquisition of H3K4me3 in heat shocked animals. **(A)** Methionine intake through diet enters the 1 carbon cycle and is used by SAM synthases for the synthesis of SAM which is used by methyltransferases to add methyl moieties to proteins, nucleic acids and lipids. **(B)** Representative confocal images of animals co-expressing RFP::SAMS-1and GFP::SAMS-4 in the germline and intestine. Kaplan-Meier survival plots of *sams-1(lof)* (**C**) or *sams-4(ok3315)* (**D**) following heat shock. Statistical significance is shown by Log-rank test. Each graph represents the compiled data from 3 biologically independent repeats; data is compiled in Table S2. Representative immunofluorescence images of intestinal nuclei stained with H3K4me3-specific antibody and quantification in *sams-1(lof)* animals (**E, H**), *sams-4(RNAi)* (**F, I**) or in *sams-1(lof); sams-4(RNAi)* animals (**G, J**). *sams-3* may also be targeted; see also **Figure S3E**). Statistical significance was calculated using unpaired Student’s t-test. ns= not significant, **** = p<0.0001, *** = p<0.001. Graph represents compiled data from three biologically independent repeats per condition.

Alterations in 1CC function can cause a variety of defects ^1^, including intriguing connections between this cycle, stress responses and aging. Lifespan lengthens in yeast, *C. elegans, Drosophila* and rodent models when methionine is restricted, genes in the methionine-SAM (Met-SAM) cycle are mutated, or polyamines are supplemented ^3^. While multiple aspects of 1CC function could affect aging, the Met-SAM cycle has particularly strong links. For example, a *C. elegans* SAM synthase, *sams-1*, was identified in a screen for long-lived animals ^4^ and multiple SAM-utilizing histone methyltransferases are also implicated as aging regulators ^5–7^. Of bioactive molecules, SAM is second only to ATP in cellular abundance ^8^, which raises the question of how such an abundant metabolite can exert specific phenotypic effects. Strikingly, studies in multiple organisms from a variety of labs have shown that reduction in SAM levels preferentially affects H3K4me3 levels ^9–12^. However, changes in SAM production may affect other histone modifications as well. For example, the Gasser lab showed that *sams-1* and *sams-3* have distinct roles in heterochromatin formation, which involves H3K9me3 ^13^ A yeast SAM synthase has also been shown to act as part of the SESAME histone modification complex ^14^ or to cooperate with the SIN3 repressor ^15^. In addition, most eukaryotes have more than one SAM synthase, which could allow partitioning of enzyme output by developmental stage, tissue type or cellular process and underlie specific phenotypic effects. Indeed, in budding yeast, SAM1 and SAM2 are co-expressed but regulated by different metabolic events, have distinct posttranslational modifications, and act differently in phenotypes such as genome stability ^16^. The two SAM synthases present in mammals are expressed in distinct tissues: MAT2A is present throughout development and in most adult tissues, whereas MAT1A is specific to adult liver ^17^. MAT2A may be present in distinct regulatory conformations with its partner MAT2B ^17^. However, the distinct molecular mechanisms impacted by these synthases are less clear. Studies exploring specificity of metazoan SAM synthase function have been difficult, as MAT1A expression decreases *ex vivo* and MAT2A is essential for cell viability ^18^. Finally, the high methionine content of traditional cell culture media has limited functional studies ^19^.

We have explored SAM synthase function in *C. elegans*, where the gene family has undergone an expansion. In *C. elegans*, genetic and molecular assays allow separation of SAM synthase expression and function *in vivo*. Furthermore, no single SAM synthase is required for survival in normal laboratory conditions or diets. *sams-1* and the highly similar *sams-3/sams-4* are expressed in adult animals, whereas *sams-5* is present at low levels in adults and *sams-2* is a pseudogene ^20^. We previously found that *sams-1* had multiple distinct functions, contributing to PC pools and stimulating lipid synthesis through a feedback loop involving *sbp-1*/SREBP-1 ^21^ as well as regulating global H3K4me3 levels in intestinal nuclei ^12^. Our studies also showed that loss of *sams-1* produced different phenotypes in bacterial or heat stress. While *sams-1* was necessary for pathogen challenge, promoter H3K4me3 and expression of immune genes, animals surprisingly survived better during heat shock when they lacked *sams-1* ^12^. Because heat shocked animals require the H3K4me3 methyltransferase *set-16/MLL* for survival, we hypothesized that SAM from a different source may be important for histone methylation and survival in the heat shock response. Here, we find that SAM source impacts the functional outputs of methylation. While the SAM and the 1CC are well associated with regulation of lifespan and stress responses, direct molecular connections have been difficult to discover. Mechanisms controlling provisioning of SAM, therefore, could provide a critical level of regulation in these processes. We show that *sams-1* and *sams-4* differentially affect different populations of histone methylation and thus gene expression in the heat shock response, and that their loss results in opposing phenotypes. Our study demonstrates that SAM synthases have a critical impact on distinct methylation targets and phenotypes associated with the stress response. Thus, defining the specificity of SAM synthases may provide a method to identify from broad effects methylation events that are specific phenotypic drivers.

## Results

### *sams-1* and *sams-4* have overlapping and distinct expression patterns and gene regulatory effects

Animals respond to stress by activating specialized protective gene expression programs ^22^. While these programs depend on specific signaling and transcriptional activators, they may also be impacted by histone methylation and the production of SAM. For example, we found that *C. elegans* lacking *sams-1* die rapidly on pathogenic bacteria, have low global H3K4me3 and fail to upregulate immune response genes ^12^. In contrast, heat shocked animals survive better without *sams-1*^23^. *sams-1(RNAi)* animals induced heat shock genes to normal levels and acquired additional changes in the transcriptome, including downregulation of many metabolic genes. However, the H3K4me3 methyltransferase *set-16*/MLL was essential for survival ^23^, suggesting that methylation was required. We hypothesized that other SAM synthases could play an important role in mediating survival during heat shock (**Fig1A**).

In order to test these hypotheses, we first compared expression of each synthase, SAM levels and gene expression after RNAi in adult unstressed animals. ModEncode data^24^ from young adult animals shows that in young adult levels, *sams-1* is expressed at the highest levels, comparable to the metabolic enzyme GAPDH (*gpdh-1*) (**FigS1A**). *sams-3* and *sams-4* are expressed at lower levels, but comparable to other enzymes of the 1-Carbon cycle such as *metr-1*, whereas *sams-5* is minimally expressed (**FigS1A**). In order to determine the tissue-specific patterns of the SAM synthases expressed in adult animals, we obtained strains where each protein was tagged with RFP, GFP or mKate, via CRISPR (**Fig1B, FigS1B, C**). RFP::*sams-1* and GFP::*sams-4* animals were also crossed to allow visualize expression of both synthases (**Fig1B**). RFP::SAMS-1fluorescence was evident in much of the adult animal, including intestine, hypodermis and cells in the head (**Fig1B, FigS1B**), in line with mRNA expression patterns derived from tissue-specific RNA seq ^25^. However, RFP::SAMS-1was not present in the germline, which did express GFP::SAMS-4 and SAMS-3::mKate (**Fig1B, S1C**). GFP::SAMS-4 and SAMS-3::mKate was also present in intestinal and hypodermal cells (**Fig1B, FigS1C**), demonstrating that these tissues, which are major contributors to the stress response ^26^ contain each of these SAM synthases. *sams-3* and *sams-4* are expressed bidirectionally from the same promoter and share 95% sequence identity at the nucleotide level thus RNAi targeting is likely to affect both genes. Indeed SAMS-3::mKate and GFP::SAMS-4 were reduced after either RNAi (**FigS1C**). Next, we used mass spectrometry to compare SAM levels after *sams-3* and *sams-4* RNAi and found that like *sams-1*^12,21^, reduction in any synthase significantly reduced but did not eliminate SAM (**Fig S1D**).

In order to compare gene expression after RNA of each SAM synthase in basal conditions, we used RNA sequencing (RNAseq). Principal component analysis showed that *sams-1(RNAi)* and *sams-5* formed distinct clusters on the first two principal components, however *sams-3* and *sams-4* were overlapping (**FigS2; Table S1: Tabs A-C**). About half of the genes upregulated after *sams-4* knockdown also increased in *sams-1(RNAi)* animals (**FigS3B)**. To determine if genes related to distinct biological processes were present, we compared genes upregulated after *sams-1* RNAi^23^ with those changing in *sams-4* RNAi with WormCat ^27^, which provides enrichment scores for three category levels (Cat1, Cat2, Cat3) for broad to more specific comparisons. WormCat finds that gene function categories at the Cat1 and Cat 2 level, such as METABOLISM: Lipid (**FigS2C**) or STRESS RESPONSE: Pathogen (**FigS2D-F**), are enriched at lower levels and contain different genes in *sams-4(RNAi)* animals (**Table S1: Tabs D-F**). Notably, *fat-7* and other lipid synthesis genes that respond to low PC in *sams-1* animals are not upregulated after *sams-4(RNAi)* (**TableS1:Tab:B**). These findings strengthen the idea that these SAM synthases could have distinct functions.

**Figure 2:**
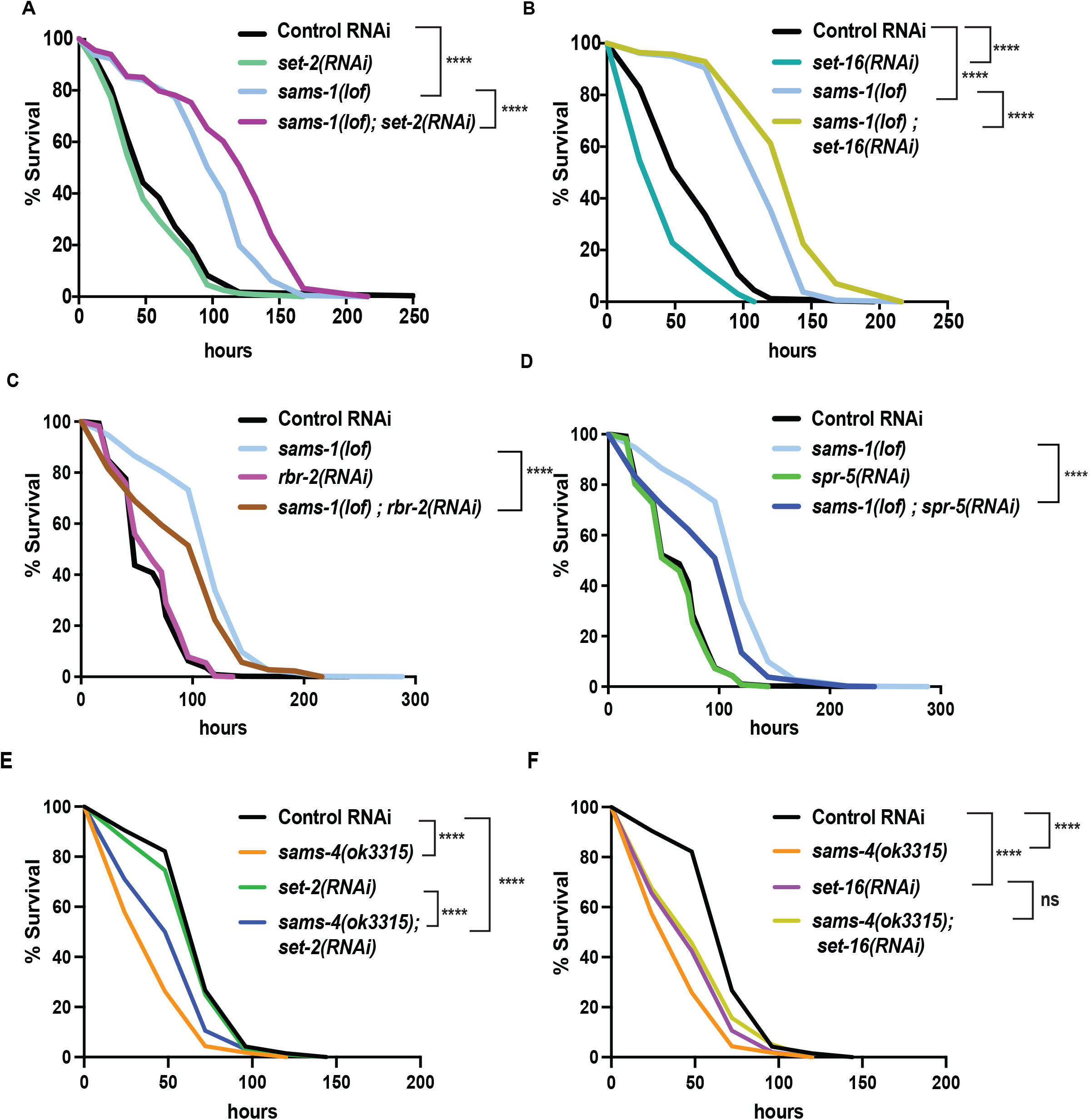
H3K4me3 demethylases modulate SAM synthase phenotypes. Kaplan-Meier plots of survival assays comparing basal and heat shocked wild type (N2) or *sams-1(lof)* animals grown on RNAi for the histone methyltransferases *set-2* (**A**) and *set-16* (**B**), or demethylases *rbr-2* (**C**) and *spr-5* (**D**). Heat shock survival assays for *sams-4(ok3315)* animals exposed to *set-2* or *set-16* RNAi are shown in **E, F**. Statistical significance is shown by Log-rank test. Each graph represents compiled data from 3 biologically independent repeats. Data for each replicate is compiled in Table S2.

### Opposing roles and requirements for *sams-1* and *sams-4* in the heat shock response

In order to determine if other SAM synthases expressed in adult animals contributed to survival in heat shock, we compared the heat shock survival phenotypes of *C. elegans* with deletions in *sams-1, sams-3* and *sams-4* to avoid effects of co-targeting by RNAi. *sams-1(ok3033)* has a deletion covering the majority of the open reading frame and extracts from these animals lack SAMS-1 protein in immunoblots^12^, therefore we refer to this allele as *sams-1(lof). sams-4(ok3315)* animals have a deletion that removes around a third of the open reading frame. Strikingly, *sams-4(ok3315)* mutants had the opposite phenotype from *sams-1(lof)*, and died rapidly after heat shock (**Fig1C, D, Table S2:Tabs B, C**). *sams-3(2932)* harbors a deletion removing most of the ORF, but in contrast to *sams-4* and *sams-1*, is indistinguishable from wild type animals in a heat shock response (**FigS3B**). Although *sams-3* may be co-targeted in RNAi experiments, we will refer solely to *sams-4* in our discussion because it has the most direct link to the heat shock phenotypes. Finally, *sams-4(RNAi)* phenotypes in the heat stress response were not linked to a general failure to thrive, as *sams-4(RNAi)* animals under basal conditions had modestly enhanced lifespan (**FigS3C; Table S2: Tab A**).

Next, we used immunostaining to compare global levels of H3K4me3 in *sams-1* and *sams-4* RNAi nuclei during heat shock. In contrast to the reduction in H3K4me3 in basal conditions in *sams-1(lof), sams-4(ok 3315)* or RNAi animals (**Fig1E-F, H-I**), we detected robust levels of H3K4me3 in *sams-1(lof)* nuclei after heat shock (2 hours at 37°C) (**Fig1E, H**), suggesting that *sams-1*-independent mechanisms act on H3K4me3 during heat shock. These increases in H3K4me3 did not appear in heat shocked *sams-4(RNAi)* intestinal nuclei (**Fig1F, I**), raising the possibility that *sams-4* contributed to the effects in *sams-1(lof)* animals. Next, we wanted to test effects of reducing both *sams-1* and *sams-4* levels on H3K4me3 during heat shock. Loss of multiple SAM synthases reduces viability in *C. elegans*^13^. In order to circumvent this, we used dietary choline to rescue PC synthesis and growth of *sams-1(RNAi)* or (*lof*) animals during development ^12,21^. *sams-1(lof); sams-4(RNAi)* animals were raised on choline until the L4 stage, then moved to normal media for 16 hours before heat shock. Immunostaining of *sams-1(lof); sams-4(RNAi)* intestines showed that *sams-4* is required for the H3K4me3 in heat shocked *sams-1(lof)* nuclei (**Fig1G, J**). These results were identical when we used RNAi to reduce *sams-1* in *sams-4(ok3315)* animals (**FigS3E**). We also asked if *sams-4* was necessary for the increased survival of *sams-1* animals after heat shock and found that the survival advantage in *sams-1(RNAi)* was decreased in *sams-4(ok3315)* animals (**FigS3D**). These results suggest that H3K4me3 may be remodeled during heat shock with SAM from distinct synthases and that *sams-4*-dependent methylation is critical for survival. Previously, it was shown that H3K4me3 deposition is independent *of sams-4* in embryonic nuclei ^13^, however, our finding that it is broadly decreased in *sams-4(RNAi)* intestinal nuclei suggests it may have important roles in H3K4 methylation in adults.

Increases in H3K4me3 have also been shown to occur in budding yeast when blocks in phospholipid synthesis relieve a drain on SAM and increase levels^28^, which we have confirmed in *C. elegans*^23^. In order to determine if SAM levels could explain differences in H3K4me3 in *sams-1* and *sams-4* animals during heat shock, we used targeted LC/MS to compare SAM, it’s precursor methionine and S-adenosylhomocysteine (SAH), the product after methyl transfer, before and after heat shock. As in our previous assays, SAM decreased significantly after *sams-1* or *sams-4(RNAi)* in basal conditions (**FigS3F**), whereas SAM levels increased in each population as *sams-1* or *sams-4* animals were shifted to 37°C for 2 hours (**FigS3F**). Levels of methionine and SAH also decreased when comparing control, *sams-1* or *sams-4(RNAi)* animals in basal vs heat shocked conditions (**FigS3G, H**), consistent with increased production and utilization of SAM. The increase in SAM in heat shocked animals is consistent with our data showing the contribution of SAMS-4 to H3K4me3 and survival in heat shocked *sams-1* animals, however, a reduction in demand for SAM if other metabolic processes are reduced after heat shock could also contribute. Finally, levels of SAM in heat shocked *sams-4(RNAi)* animals also rise to levels comparable to control animals at basal temperatures, however, H3K4me3 remains low in these conditions.

### Histone methyltransferase and histone demethylation machinery have modest, separable effects on *sams* mutant heat shock phenotypes

SAM is necessary for histone methylation; however, histone methylation dynamics are also influenced by methyltransferase (KMT) or demethylase (KDMT) activity ^29^. Therefore, changes in histone methylation dynamics could also impact H3K4me3 patterns during heat shock. H3K4me3 is catalyzed by multiple versions of the COMPASS complex, which each consist of one of several SET domain histone methyltransferases and several shared accessory subunits ^30^. In mammals, seven methyltransferases in the SET1, MLL or THX groups can methylate H3K4. *C. elegans* contain single orthologs from two of these groups: *set-2/* SET1 and *set-16/MLL*,respectively, with roles in embryonic development ^31–33^, lipid accumulation and transgenerational inheritance ^6,7^. In adult *C. elegans, set-2* RNAi results in extensive loss of H3K4me3 in intestinal nuclei and although *set-16(RNAi)* causes an intermediate reduction in bulk H3K4me3 levels, it has a broader requirement for survival during stress ^23^. Because specificity for H3K4 mono, di or trimethylation has not been verified on a genome-wide scale for KDMTs, we examined multiple members of the H3K4 KDM family.

In order to determine if KMTs or KDMT dynamics played a role in the change of H3K4me3 during heat shock, we used RNAi to deplete them in *sams-1(lof)* or *sams-4(ok3315)* animals and measured survival after heat shock and intestinal H3K4me3 levels. RNAi of *set-2/SET1* (**Fig2A**) or *set-16/MLL* (**Fig2B**) increased survival in *sams-1(lof)* animals after heat shock (also **Table S2:Tabs:C, E**) and did not limit heat shock-induced H3K4me3 in *sams-1(RNAi)* nuclei (**FigS4A, B**). RNAi of two KDMTs, *rbr-2* (**Fig2C**) and *spr-5* (**Fig2D**) had opposite effects from the KMTs, moderately reducing survival (**TableS2: Tab F**), whereas *amx-1* and *lsd-1* had no effect (**FigS4G, H; TableS2: Tabs I, J**). RNAi of *set-2* (**Fig2E**) or *set-16* (**Fig2F**) had slight, but statistically significant effects, increasing survival of *sams-4(ok3315)* animals (**TableS2: Tabs G, H**). However, survival was still significantly below controls in *sams-4(ok3315)* with or without the KMT RNAi. Taken together, this suggests that *set-2* and *set-16* may act redundantly in the deposition of H3K4me3 after heat shock and are important to survival in *sams-1(lof)* animals. Furthermore, our data illustrate that the context is critical for understanding role of SAM and H3K4me3 in stress; *sams-4* and *set-16* are generally required for survival after heat shock, but loss of either H3K4 KTM enhances survival in *sams-1(lof)* animals.

### Distinct patterns of H3K4me3 and gene expression in *sams-1(RNAi)* versus *sams-4(RNAi)* animals during heat shock

H3K4me3 is a prevalent modification enriched near the transcription start sites (TSSs) of actively expressed genes ^34^. Differing global patterns of H3K4me3 in *sams-1(RNAi)* and *sams-4(RNAi)* nuclei suggest this histone modification at specific sites could also be distinct. In order to identify loci that might link H3K4me3 to these phenotypes, we used CUT&Tag, (Cleavage Under Targets and Tagmentation, C&T) ^35^, to determine genome-wide H3K4me3 levels in Control RNAi, *sams-1* and *sams-4(RNAi)* in basal (15°C) and after heat shock (37°C/2 hours) from two biologically independent replicates along with no antibody controls. C&T is uniquely suited to the small sample sizes available from these stressed populations. In this approach, a proteinA-Tn5 transposase fusion protein binds to the target antibody in native chromatin and DNA libraries corresponding to antibody binding sites are generated after transposase activation. After sequencing of libraries, we used the HOMER analysis suite^36^ to analyze reads mapped to the *C. elegans* genome and called peaks using ChIPSeqAnno^37^ for more detailed peak annotation. Bar plots from ChIPSeqAnno annotations and TSS plots generated with HOMER show robust mapping of H3K4me3 to promoter-TSS regions, validating this approach (**Fig3A; TableS3: Tabs A-F**). While promoter-TSS regions were the largest feature in each sample, heat shocked *sams-4(RNAi)* animals had fewer overall peaks (**Fig3A**). Correlation plots also show strong similarity between replicates (**FigS5A**). Because C&T has not been extensively used in *C. elegans*, we compared data from basal conditions in our study to three previously published ChIP-Seq data sets^38,39,40^. We compared our C&T data from wild type young adult animals grown at 15°C on control RNAi food (HT115) against ModEncode (L3 animals), *glp-1(e2141)* mutants from Pu et al.^41^ and wild type adults grown at 20°C on OP50 bacteria from Wan et al. ^40^ by computing a pair-wise Pearson correlation. We found our C&T clustered most closely with the ChiPSeq from wild type animals in Wan et al., along with one of the modEndode replicates (**FigS5B**) with moderate correlation scores. Both our C&T data and the Wan ChiPseq data correlated poorly with the Pu et al. ChIP seq, which is likely due to the lack of germline nuclei in these animals. The moderate correlation between our data and ChiP seq from Wan et al may be due to differences in growth temperature and bacterial diet. As a part of our quality control, we visually inspected browser tracks around the *pcaf-1* gene, which is a long gene and has been used by our labs and others as a positive control for H3K4me3 localization in the 5 prime regions ^12,32^. H3K4me3 peaks are prominent upstream of the transcript as expected and the no antibody libraries showed few reads (**FigS5C**).

**Figure 3.**
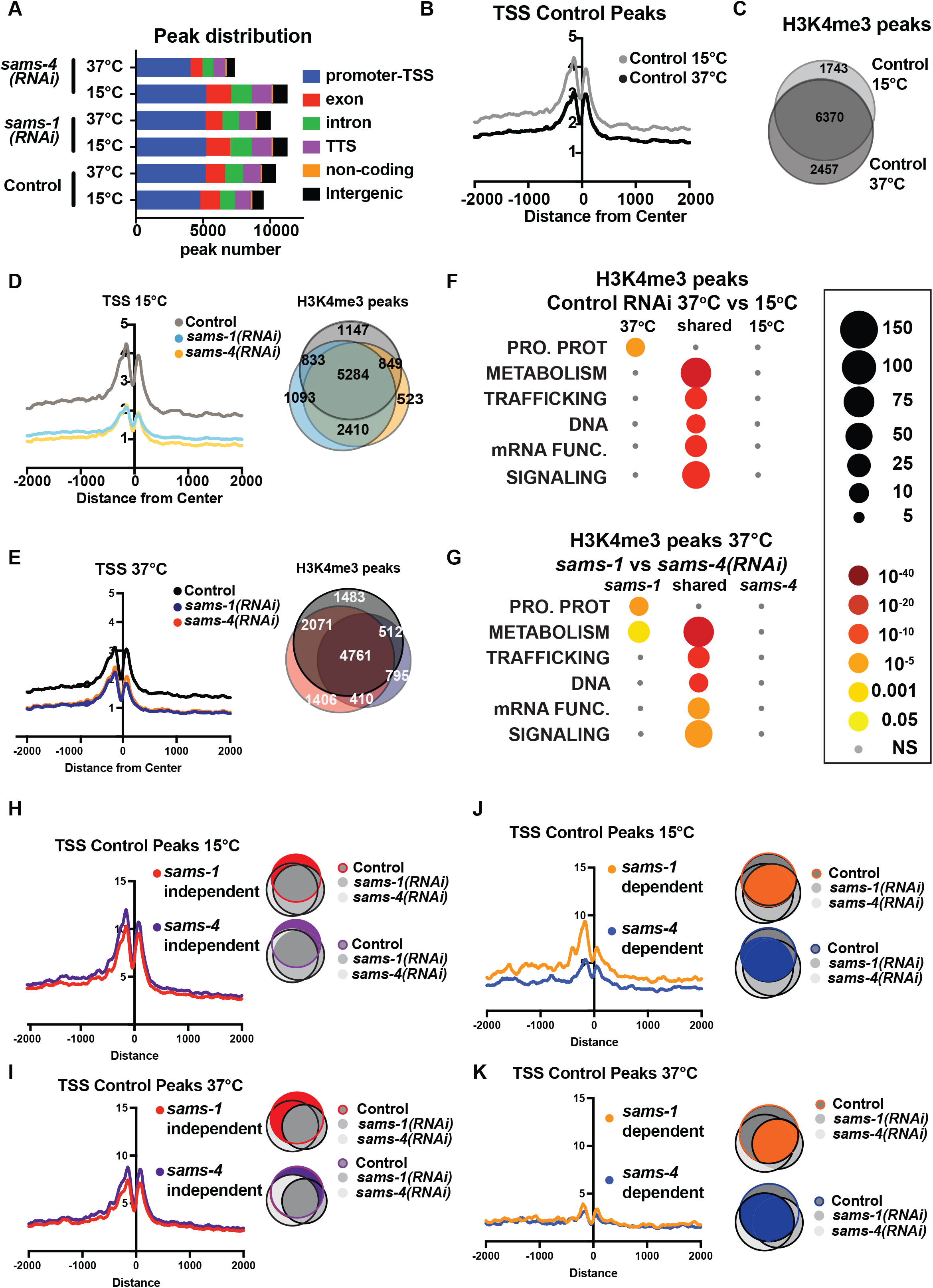
H3K4me3 modifying enzymes modulate SAM synthase phenotypes. **(A)** Bar graph showing the distribution of the enrichment of H3K4me3 over different genomic loci in animals fed control RNAi, *sams-1(RNAi)* or *sams-4(RNAi)* at 15°C and 37°C. **(B)** Aggregation plots showing TSS enrichment in the H3K4me3 peaks identified in animals fed control RNAi at 15°C and 37°C. The Y axis on TSS plots shows Peaks per base pair of gene. **(C)** Venn diagram comparing the overlap in the H3K4me3 peaks identified in animals fed control RNAi at 15°C and 37°C. **(D)** Aggregation plots showing TSS enrichment in the H3K4me3 peaks identified in animals fed control RNAi or *sams-1(RNAi)* or *sams-4(RNAi)* at 15°C and Venn diagram comparing the overlap in the H3K4me3 peaks identified in animals fed control RNAi or *sams-1(RNAi)* or *sams-4(RNAi)* at 15°C. **(E)** Aggregation plots showing TSS enrichment in the H3K4me3 peaks identified in animals fed control RNAi or *sams-1(RNAi)* or *sams-4(RNAi)* at 15°C and Venn diagram comparing the overlap in the H3K4me3 peaks identified in animals fed control RNAi or *sams-1(RNAi)* or *sams-4(RNAi)* at 37°C. **(F)** Bubble chart showing enriched gene categories in differential peaks as determined by WormCat in animals fed control RNAi at 15°C only, 37°C only and common between 15°C and 37°C (**G**) or *sams-1(RNAi)* and *sams-4(RNAi)* at 37°C. Aggregation plots showing TSS enrichment of Control peaks that did not change after *sams-1(RNAi)* and *sams-4(RNAi)*(independent) **(H)** 15°C or **(I)** 37°C. Shaded areas in the Venn diagrams indicate the population of genes used for plotting the TSS enrichment plots. Aggregation plots showing TSS enrichment of Control peaks that were dependent on *sams-1(RNAi)* or *sams-4(RNAi)* **(J)** 15°C or **(K)** 37°C. Shaded areas in the Venn diagrams indicate the population of genes used for plotting the TSS enrichment plots.

Next, we compared TSS distributions and examined overlap between H3K4me3 peaks in Control RNAi animals in basal and heat shock conditions and found moderate reductions occurred with heat shock (**Fig3B**). Around 20-30% of peaks were specific to at either at basal (15°C) vs. heat shock (37°C) temperature (**Fig3C)**, suggesting that H3K4me3 could be remodeled upon heat shock in *C. elegans*. TSS enrichment of H3K4me3 was sharply reduced in both *sams-1* and *sams-4* samples at 15°C, however this difference was less marked in heat shocked animals, in line with lower TSS localization in Control animals (**Fig3D, E**). While aggregate TSS enrichment for H3K4me3 was similar for *sams-1* and *sams-4* RNAi animals, this analysis could miss distinct sets of H3K4me3 marked genes in each condition. Indeed, Control, *sams-1* and *sams-4(RNAi)* animals each showed 500-1000 specific peaks in basal conditions, with moderate increases in these numbers after heat shock (**Fig3D, E**). As H3K4me3 is a widely occurring modification, we hypothesized that we might better understand potential SAM synthase-specific requirements if we focused on peaks that change in the Control RNAi heat shock response and asked how they are affected by loss of *sams-1* or *sams-4*. We used two different methods for comparing potential SAM synthase requirements for H3K4me3 in the heat shock response. First, we used differential peak calling (ChIPPeakAnno ^37^) followed by WormCat category enrichment to determine the classes of genes which might be affected (**FigS6A-F; TableS3; Tabs G-I**). Peaks present in both basal and heat shocked conditions were enriched for genes in the METABOLISM category (including Lipid: phospholipid, sphingolipid, sterol and lipid binding, along with mitochondrial genes) as well as in core function categories such as those involved in trafficking, DNA or mRNA functions (**Fig3F, FigS6D-E; Table S3: Tabs G-I)**. There was no significant category enrichment specific to 15°C animals, but after heat shock, Control RNAi animals gain enrichment in peaks at the Category 1 level in PROTEOSOME PROTEOLYSIS (**Fig3F**). This enrichment was driven by increases in H3K4me3 at E3: Fbox genes (**FigS6A, B; Table S3:Tab A,B**), which could be important for eliminating mis-folded proteins during heat shock. Comparison of peaks differentially present in *sams-1* and *sams-4* RNAi animals showed that only *sams-1(RNAi)* exhibited a similar enrichment to Control RNAi in the PROTEOLYSIS PROTEOSOME category (**Fig3G, FigS6C, D**), which could help explain the reduced survival of *sams-4(RNAi)* animals relative to *sams-1(RNAi)* animals. *sams-1* RNAi animals also gained enriched peaks in a wide range of gene categories within METABOLISM, whereas *sams-4(RNAi)* enriched peaks in these categories were more limited (**FigS6C-F**). Thus, loss of *sams-1* or *sams-4* differentially affects H3K4me3 peaks within functional gene classes that also change in the heat shock response.

Next, we hypothesized that H3K4me3 at peaks in Control RNAi animals might reflect multiple differently regulated populations, some which are linked to SAM synthase function and others that are regulated at other levels. In order to test this, we divided peaks in Control animals at 15°C or 37°C into those that did not change after SAM synthase RNAi (*sams-1* or *sams-4* independent peaks) or those that were dependent on *sams-1* or *sams-4* and examined aggregations around TSS regions. There was little difference between TSS plots of *sams-1* or *sams-4*-independent genes at either temperature (**Fig3H, I**). However, in basal conditions, Control peaks that depended on *sams-1* had more marked TSS localization (**Fig3J**), demonstrating that *sams-1* and *sams-4* dependent peaks have distinct TSS architectures. TSS localization was low in all 37°C samples, following the general trend of decrease after heat shock (**Fig3K**). We next separated Control peaks into those that were generally SAM synthase-dependent and those that were specific to loss of *sams-1* or *sams-4*. Aggregation of these peaks shows that peaks in Control 15°C samples that were lost only in *sams-4* RNAi also had the lowest levels of H3K4me3 in TSS regions, whereas promoters that lost this modification only after *sams-1* RNAi had higher levels of H3K4me3 (**FigS6G**). Control 37°C samples exhibited a similar pattern, with a lower H3K4me3 level overall consistent with what we have observed in heat shock samples (**FigS6H**). Thus, genome wide H3K4me3 contain multiple populations with distinct TSS patterns. Peaks that are present even when *sams-1* or *sams-4* are depleted have the highest levels, whereas *sams-1*-dependent peaks have moderate H3K4me3, and peaks that are lost after *sams-4* RNAi have the lowest levels. Taken together, this shows that individual SAM synthases are linked to distinct sets of H3K4me3 within the genome.

### RNAi of *sams-1* or *sams-4* has similar effects on TSS peaks at tissue-specific genes

Our C&T and RNA seq assays were performed on whole animals. While *sams-1* and *sams-4* are co-expressed in the intestine and hypodermis, which are major stress-responsive tissues, the germline nuclei contain only *sams-4* (**Fig1B** and **FigS1B**). This aligns with our previous observations that *sams-1(RNAi)* animals had normal patterns of H3K4me3 in germline nuclei (Ding, et al. 2015), whereas RNAi of *sams-4* abrogates H3K4me3 staining in germline nuclei (**FigS7A**). However, embryo production and development appear broadly normal in *sams-4* RNAi embryos (not shown). In order to determine how H3K4me3 might align with tissue-specific expression patterns, we aggregated peaks from tissue-specific RNA seq data published by Serizay, et al ^42^. Serizay et al. separated nuclei based on tissue specific GFP expression and defined gene sets that were expressed that were ubiquitously, as well as those that were present only in a single tissue. They also performed ATAC seq (Assay for Transposase-Accessible Chromatin using sequencing). Serizay, et al. defined transcripts by expression pattern and defined sets that were specific to (*tissue*_only), or represented in across multiple tissues (*tissue_all*). ubiquitious_all and Germline_only genes had the most defined patterns of open chromatin around TSSs ^42^ We compared our C&T data with Ubiquitious_all, Germline_only and Intestine_only genes and found that we identified peaks for around half of these genes in Control RNAi animals at 15°C or 37°C (**FigS7B-D)**. We found the ubiquitous_all and germline_only genes also had strong H3K4me3 peaks that were reduced equally by *sams-1* or *sams-4* RNAi in both temperature conditions (**FigS7D, G; E, H**). Intestine_only genes showed lower TSS enrichment but were similarly reduced after *sams-1* or *sams-4* RNAi (**FigS7H, I**). These data suggest that differences in germline expression for *sams-1* and *sams-4* are not sufficient to explain differential effects on H3K4me3 peak populations.

### Poor expression of heat shock gene suite in *sams-4(RNAi)* animals

H3K4me3 is found at the promoters of many actively transcribed genes, but it is not necessarily required for gene expression ^29^. However, studying chromatin modification in stress responses may reveal additional regulatory effects ^43^. We previously found using ChIP-PCR in the context of the stress response in *C. elegans* that H3K4me3 increased at promoters of genes that responded to bacterial stress in a *sams-1-* dependent manner^12^. However, during the stress response, H3K4me3 did not change at multiple non-stress responsive genes, suggesting that stress-responsive loci might be more sensitive to SAM levels^12^. In order to identify genes that changed in SAM-deficient animals, we performed RNA seq, then compared genes induced by heat shock in control and *sams-1(RNAi)*^23^ with genes induced in *sams-4(RNAi)* animals **(TableS4: A-C**). Upregulated genes for control and *sams-1(RNAi)* animals appeared closely grouped in principal component analysis, with sams-4(RNAi) upregulated genes and all downregulated gene sets forming distinct groups (**FigS8A**). We previously noted that while *sams-1(RNAi)* animals could not mount the full transcriptional response to bacterial stress, most genes activated by heat increased similarly to controls^23^. *sams-4(RNAi)* animals, in contrast, activate less than 25% of the genes induced by heat in control animals (**Fig4A**). *sams-1(RNAi)* and *sams-4(RNAi)* animals also induce more that 600 genes in response to heat that are SAM-synthase-specific and which do not increase in control animals (**Fig4A**). WormCat pathway analysis shows that *sams-4(RNAi)* animals lack the robust enrichment in STRESS RESPONSE (Cat1) and STRESS RESPONSE: Heat (Cat2) evidenced in both Control and *sams-1(RNAi)* samples (**Fig4B; TableS4: D-F**). In addition, enrichment of the CHAPERONE, PROTEOLYSIS PROTEOSOME categories occurring in *sams-1(RNAi)* animals does not occur after *sams-4(RNAi)*, reflecting lack of induction of these genes which could be important for proteostasis in the heat shock response (**Fig4C**). Thus, reduction in *sams-1* or *sams-4* results in distinct gene expression programs in both basal conditions (**FigS2A-F**) and during the heat stress response (**Fig4A-C**). This differentiation of gene expression programs clearly shows that *sams-1* and *sams-4* have distinct functional roles.

**Figure 4.**
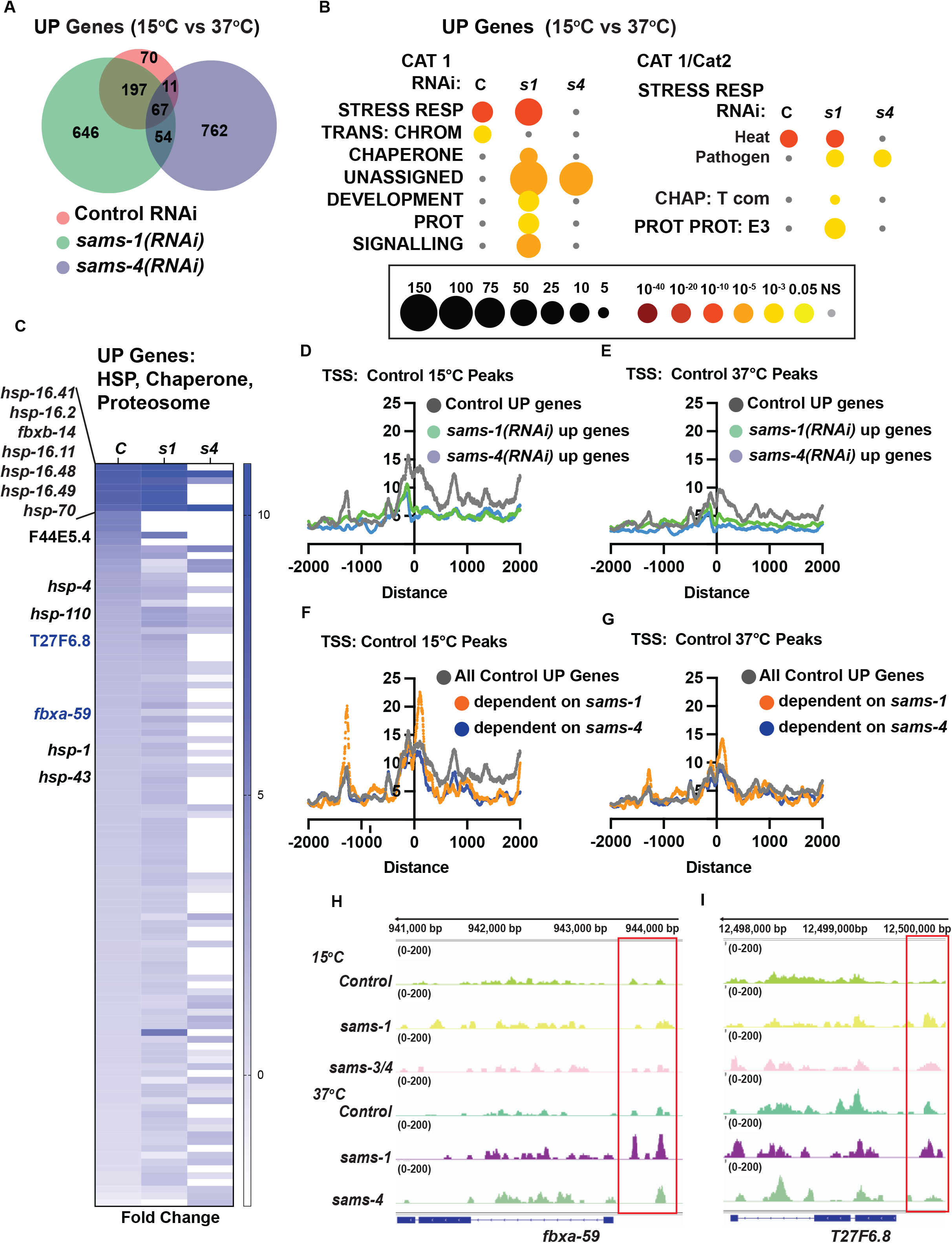
Distinct gene expression and H3K4me3 patterns after heat shock in *sams-1* and *sams-4* RNAi animals. **(A)** Venn diagram showing overlap of genes upregulated by heat shock in control, *sams-1* or *sams-4* RNAi animals. *sams-1* data is from Ding, et al. 2018. **(B)** Bubble charts show broad category enrichment of up genes determined by Worm-Cat in control (RNAi) or *sams-1* or *sams-4* animals in genes changed (FDR<0.01) after heat shock. **(C)** Heat map for heat shock response genes up regulated following heat shock in animals fed control RNAi, *sams-1* or *sams-4(RNAi)*. TSS plots showing aggregation of H3K4me3 in genes upregulated in control, *sams-1* or *sams-4* RNAi at **(D)** 15°C or **(E)** 37°C. TSS plots showing aggregation of H3K4me3 in all genes upregulated in control or *sams-1* dependent or *sams-4* RNAi dependent at **(F)** 15°C or **(G)** 37°C. The Y axis on TSS plots shows Peaks per base pair of gene. Genome browser tracks for **(H)** *fbxa-59* and **(I)** *T27F6.8* to visualize changes in H3K4me3 enrichment in animals fed control, *sams-1* or *sams-4(RNAi)* at 15°C or 37°C.

Gene expression changes occurring after *sams-1* or *sams-4* depletion could result from direct effects on H3K4me3 or other potential methylation targets, or from indirect effects. Evaluating the impact H3K4me3 on gene expression is also complex, as this modification is generally associated but not necessary for expression of actively transcribed genes ^29^. In our analysis of H3K4me3 peaks during the heat stress response, we found evidence of multiple peak populations that depend on or occur independently of *sams-1* or *sams-4* (**Fig2H-K, FigS6A-F**). We reasoned, therefore, that it was also critical to determine H3K4me3 levels at *sams-1-* or *sams-4*-dependent genes in the heat shock response.

First, we examined H3K4me3 peak levels at genes with increased in Control RNAi, *sams-1(RNAi)* or *sams-4(RNAi)* during heat shock. We found that genes dependent on *sams-1* or *sams-4* in the heat shock response were marked by lower overall H3K4me3 levels at the TSSs (**Fig4D**). However, this analysis included large numbers of upregulated genes in *sams-1* or *sams-4* outside of the wildtype heat stress response.

Therefore, we next focused on genes normally upregulated during heat shock and divided them according to SAM synthase dependence. Strikingly, isolating the *sams-1-* dependent genes revealed a strong peak 5’ to the TSS, which was not evident in the larger subset of Control or *sams-4(RNAi)-dependent* upregulated genes (**Fig4E, F**). Among the genes with robust peaks in heat shocked *sams-1(RNAi)* animals were two F-box proteins, *fbxa-59* and T27F6.8, which were robustly expressed in *sams-1* but not *sams-4* animals (**Fig4C, G-I**). Down regulation of T27F6.8 did not affect the survival of the animals after heat shock (**FigS8B**) while survival of animals fed *fbxa-59* RNAi was modestly affected (**FigS8C**). Survival in heat shock may be multi-genic and rely on pathway responses rather than single genes. However, our data reveals genes upregulated in the heat shock response may have different H3K4me3 levels depending on requirements for *sams-1* or *sams-4*. In addition, our results suggest that roles for H3K4me3 may become clearer when genome-wide methylation populations are separated into biologically responsive categories.

### SAM synthase-specific effects on genes downregulated in the heat shock response

Transcriptional responses to heat shock largely focus on rapidly induced genes that provide protection from changes in proteostasis ^44,45^. However, downregulated genes could also play important roles. For example, the WormCat category of TRANSMEMBRANE TRANSPORT (TM TRANS) is enriched in genes downregulated after heat shock in *C. elegans* (**Fig5A, B**). Previously we observed that heat shocked animals depended on *sams-1* for normal expression of nearly 2,000 genes, falling within WormCat Categories of METABOLISM, TRANSCRIPTION FACTOR (TF), SIGNALLING and STRESS RESPONSE^23^ (**Fig5A, B**). Interestingly, the metabolic genes dependent on *sams-1* include those in lipid metabolism, whereas the TF enrichment was centered around nuclear hormone receptors (NHRs) (**Fig5C, D**), which regulate many metabolic and stress responsive genes in *C. elegans*^46^. However, neither the shared TM TRANSPORT nor the *sams-1* specific categories depend on *sams-4* (**Fig5B, C**). Thus, as in genes upregulated during the heat shock response, genes downregulated in the heat shock response also have differential requirements for *sams-1* and *sams-4*.

**Figure 5.**
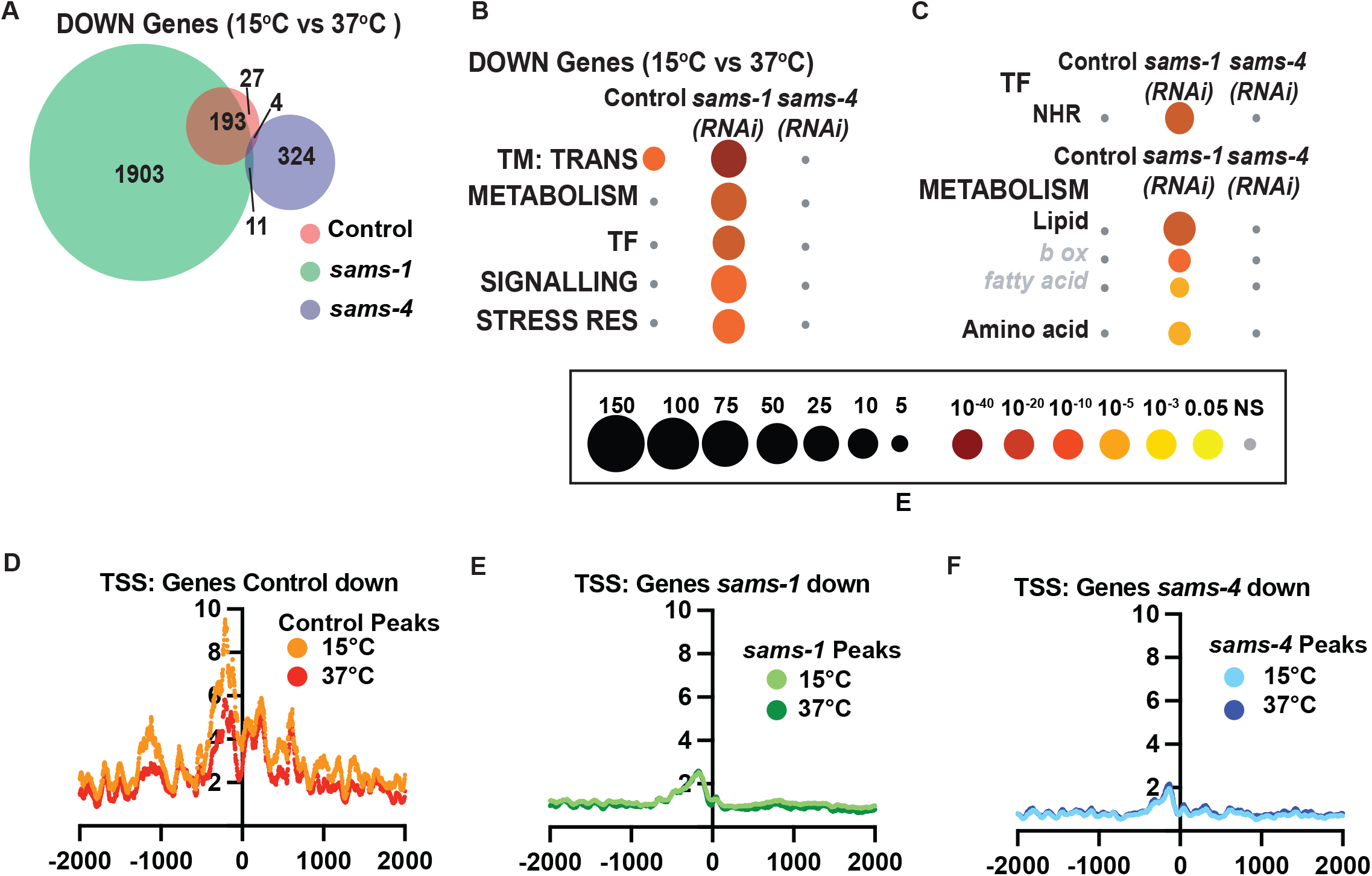
Genes that depend on *sams-1* or *sams-4* for expression have reduced H3K4me3. **(A)** Venn diagram showing overlap in down regulated genes in animals fed control, *sams-1* or *sams-4(RNAi)* at 37°C. **(B)** Bubble charts show broad category enrichment of metabolism genes determined by Worm-Cat in *sams-1* or *sams-4* animals in genes changed (FDR<0.01) after heat shock. **(C)** Bubble charts show broad category enrichment of transcription factor and metabolism genes determined by Worm-Cat in *sams-1* or *sams-4* animals in genes changed (FDR<0.01) after heat shock. Aggregation plots showing average enrichment of reads around the transcription start site (TSS) in animals fed **(D)** control, **(E)** *sams-1* or **(F)** *sams-4(RNAi)* at 15°C or 37°C. The Y axis on TSS plots shows Peaks per base pair of gene.

Next, we examined H3K4me3 levels around TSSs of genes that lost expression during heat shock in Control, *sams-1* or *sams-4(RNAi)* animals. Genes decreasing in Control animals had a slight reduction of H3K4me3 peaks when comparing15°C and 37°C samples, consistent with global levels after heat shock (**Fig5D**). RNAi of *sams-1 or sams-4* also broadly reduced H3K4me3 TSS enrichment at downregulated genes (**Fig5D-F**). However, there were minimal differences before or after heat shock, suggesting expression patterns affecting survival could be established before induction of the stress.

H3K4me3 has been reported to act as a bookmarking modification, therefore we hypothesized that some loci could be affected before heat shock, with expression changing afterward. Therefore, we more closely examined genes with *sams-1-* dependent H3K4me3 at 15°C that lost expression during heat shock. Those genes were highly enriched for METABOLISM: Lipid: beta oxidation and NHR transcription factors (**Fig6A**). We noted they included multiple members of a regulatory circuit that control expression of a beta-oxidation-like pathway that degrades toxic fatty acids identified by the Walhout lab ^47^, including *nhr-68, nhr-114* and beta-oxidation genes *acdh-1, hach-1 ech-6, −8*, and *-9* **(Fig6B, C**). Indeed, *nhr-68*, the initiating TF in this regulatory circuit, shows lower levels of H3K4me3 at its promoter in basal conditions, compared to Control or *sams-4* RNAi animals (**Fig6D**). The H3K4me3 peak overlaps with another gene, *pms-2*, whose expression does not change after heat shock or upon SAM synthase RNAi (**TableS4: Tabs A-C**). In order to test if H3K4me3-dependent regulation of *nhr-68* was important for survival during heat shock, we made use of a construct expressing *nhr-68* under the intestine-specific *ges-1* promoter^47^, where H3K4me3 peaks do not change after RNAi of *sams-1* or *sams-4* (**FigS9A**). Expression of *nhr-68* under this heterologous promoter had a moderate, but significant effect on survival (**Fig6E**). Thus, downregulation of *nhr-68* in sams-1 animals after heat shock could be part of a program enhancing survival. Taken together, our results suggest differences in H3K4me3 patterns in *sams-1* and *sams-4* animals before heat shock may also influence gene expression patterns during the stress response. This demonstrates that *sams-1* and *sams-4* are required for distinct sets of genes in the heat stress response and contribute to different H3K4me3 patterns.

**Figure 6.**
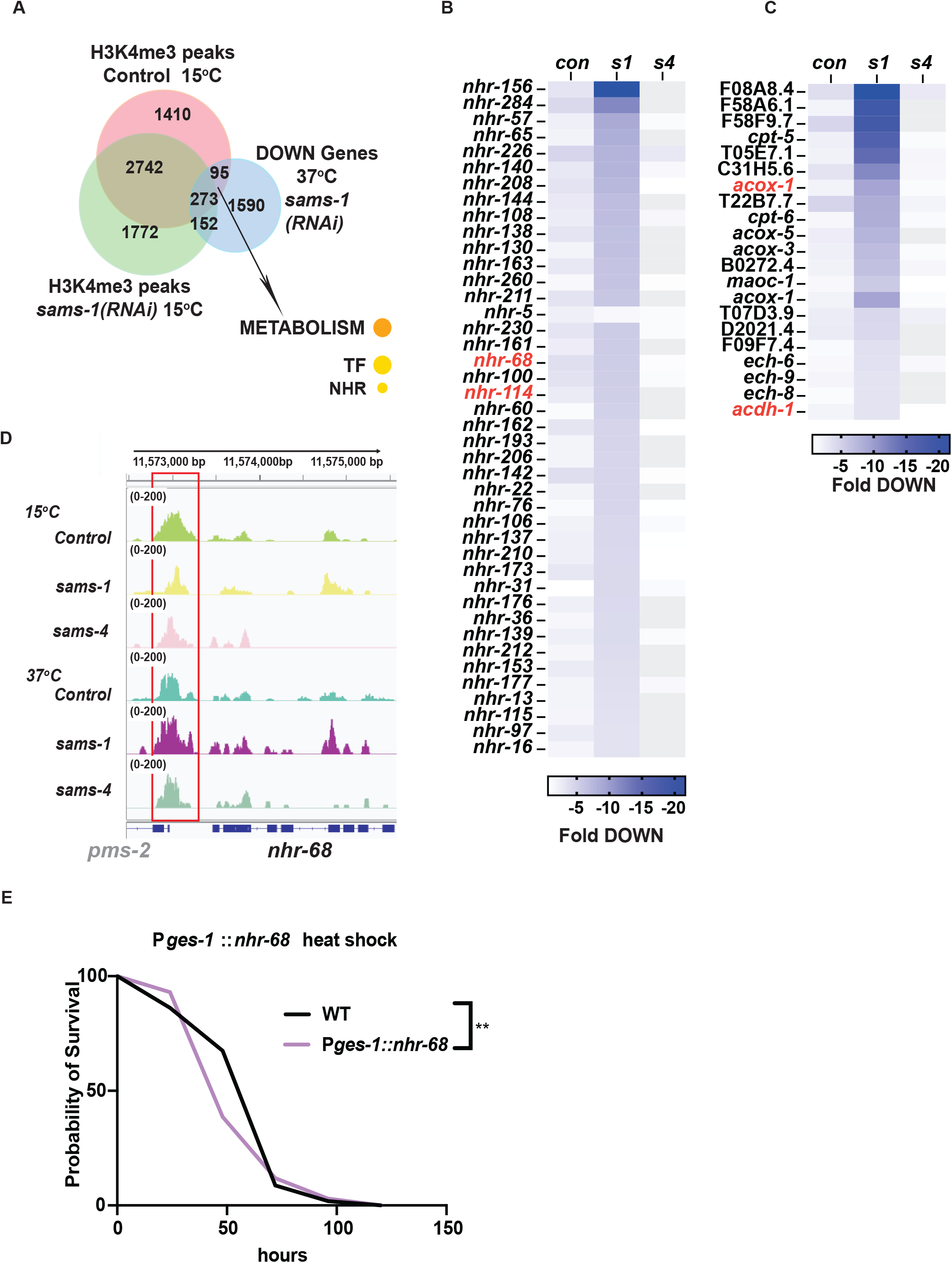
*nhr* and lipid beta oxidation genes lose H3K4me3 after *sams-1* RNAi but expression after heat shock. (**A**) Venn diagram showing the overlap between H3K4me3 peaks identified in animals fed control or *sams-1(RNAi)* at 15°C and down regulated genes identified in heat shocked animals fed *sams-1(RNAi)*. Heat map for **(B)** nuclear hormone response genes and **(C)** lipid β-oxidation genes down regulated following heat shock in animals fed control RNAi, *sams-1* or *sams-4(RNAi)*. Genes linked to nhr-68 feedback loop^47^ are marked in red. **(D)** Genome browser tracks for *nhr-68* to visualize changes in H3K4me3 enrichment in animals fed control, *sams-1* or *sams-4(RNAi)* at 15°C or 37°C.

## Discussion

The molecules that modify chromatin are produced by metabolic pathways ^48^. Use of ATP, AcetylCoA or SAM for phosphorylation, acetylation or methylation of histones is tightly regulated and many studies have focused on control of enzymes or enzyme-containing complexes. Acetylation and methylation may also be regulated by metabolite levels ^49,50^. This allows the chromatin environment to sense and respond to changes in key metabolic pathways. However, effects of methylation on chromatin are multifaceted: DNA and H3K9me9 have strong repressive effects, whereas other modifications such as H3K4me3 and H3K36me3 are associated with active transcription ^29^. These marks, especially H3K4me3, are most sensitive to SAM levels, most likely due to the kinetics of the H3K4me3 MTs ^51^. SAM is an abundant metabolite that contributes to multiple biosynthetic pathways in addition to acting as the major donor for histone, DNA and RNA methylation ^52^. Reduction in SAM levels has major phenotypic consequences in animals, altering lipid levels in murine liver and *C. elegans*^21,53^, altering differentiation potential in iPS cells ^10^ and changing stress resistance ^23^. In addition, 1CC has been identified as a causal regulator of aging ^54^ and is important in cancer development ^19,55^. However, the abundance of SAM and its targets have made it difficult to connect changes in methylation to molecular pathways regulating these physiological effects. In addition, studying effects of SAM is difficult in culture because SAM itself is labile ^56^ and tissue culture media is replete with 1CC metabolites ^19^. Important insights have been made into the impact of SAM on the breadth of H3K4me3 peaks using methionine depletion ^9,57,58^, however, this approach could affect other pathways. In this study, we have taken the approach of limiting SAM synthase expression in *C. elegans*, then using genetic and molecular approaches to link methylation-dependent pathways to changes in stress responses. We found that individual SAM synthases could have distinct effects even on a single methylation target such as H3K4me3. This observation not only shows that examining how SAM is produced within the cells allows differentiation of phenotypic effects, but also supports the striking notion of ‘where’ SAM comes from affects its functional output. While mammalian cells express either one of two SAM synthases, MAT2A, which is present in non-liver cells, may be present in multiple regulatory isoforms ^59^. Thus, the isoform-specific production and functional targets for SAM synthases we uncover could also be important in mammals. Hints of this exist in other cellular systems – 1CC enzymes, for example, have been associated with chromatin modifying complex in yeast ^14^ and mammalian cells ^60^.

H3K4me3 is clearly an important link between SAM levels, aging and stress phenotypes, as loss or reduction of H3K4 MT function phenocopy aspects of SAM depletion ^12,23^. However, this modification is also wide-spread, and transcription may occur even when this mark is not present ^61^. By studying acute changes in gene expression during heat stress response in *C. elegans*, we have found that H3K4me3 populations can also be separated based on SAM synthase requirements. The importance of H3K4me3 during heat shock is also reflected in the interactions between the SAM synthases and the KMTs/KDMTs as lowering levels of *set-2* or *set-16* increase survival. This suggests that the context of low SAM from SAMS-1, reducing H3K4me3 can have additional benefits. Future studies identifying genomic targets of H3K4me3 KMTs together with SAM synthases may be important for untangling these effects.

SAM synthase-specific effects may also vary according to the biological context, as loss of *sams-1* improves the ability of *C. elegans* to survive heat stress, while limiting its ability to withstand bacterial pathogens ^23^. Our previous studies showed that the induction of bacterial pathogen induced genes was limited in the absence of *sams-1*,however, in this study, we find links between *sams-1*-dependent genes in basal conditions and effects on survival after heat shock. Thus, the altered methylation landscape in *sams-1* animals provides a context favorable to extended lifespan and survival in heat stress but which limits other stress responsive genes. This context may depend on systems level effects and not on a single “target” gene, as our analysis of genes that lose peaks in *sams-1* or *sams-4(RNAi)* animals have modest effects, but do not recapitulate the entire phenotype. It is also possible that there are genes or specific modules that drive enhanced survival in *sams-1* animal or responsible for viability after loss of *sams-4(RNAi)*. Our approach dividing peaks into groups based on responsiveness to *sams-1* or *sams-4* demonstrates the importance of identifying specific populations of H3K4me3; combining *set-2* or *set-16* sensitive loci may provide the resolution to identify these loci in future studies. Manipulation of the 1CC is of interest as a modulator of aging^54^ and affects multiple biological processes. Our studies demonstrate that lowering SAM, or reducing levels of a key methylation target such as H3K4me3, does not represent a single biological state and that it is important to consider that effects may depend on synthase-specific regulation or context. Future identification of these regulators will provide the mechanistic details key to understanding the role of the 1CC in aging and stress.

## Limitations

The genetic tools used in our study provide a method to reduce SAM from a specific enzymatic source. However, the roles for SAM in the cell are broad and can affect methylation of multiple targets. While our metabolomics assays show that SAM increases in heat shocked *sams-1(RNAi)* animals, we have not demonstrated that this SAM is derived from *sams-4*. In addition, survival benefits after heat shock occur across broad cellular functions including proteostasis and other methylation marks such as H3K9me3 ^62^. Thus, there may be multiple additional methylation-dependent mechanisms that influence survival of *sams-1* or *sams-4* animals during heat shock. In addition, we measured gene expression and H3K4me3 at two hours post heat shock, whereas the survival assay occurs over multiple days. Thus, there may be changes in gene expression or histone modifications occurring at later times that also affect survival.

## Materials and methods

### C. elegans strains

N2(Caenorhabditis Genetics Center); *sams-1(lof)(ok3033); sams-3(ok2932) IV, sams-4((ok3315)* IV, Caenorhabditis Genetics Center), tagRFP::SAMS-1 (WAL500, this study); GFP::SAM-4(WALK501, this study); SAMS-1::RFP;GFP::SAMS-4(WAL502, this study), SAMS-3::mKate (WAL305). *Pges-1*::NHR-68::GFP (VL1296)^47^. CRISPR tagging for WAL500 and WAL501 were done by the UMASS Medical School transgenic core, confirmed by PCR for genotype and outcrossed three times to wild type animals. Next, each strain was crossed to the respective deletion allele to create WAL503 (*RFP::sams-1(ker5); sams-1(ok3033)*) and *WAL504(GFP::sams-4(ker6); sams-4(ok3315)). sams-3::mKate(nu3139)* (COP2476) was constructed using CRISPR by In Vivo biosystems then outcrossed 3 times (WAL305).

### *C. elegans* culture, RNAi and stress applications

*C. elegans* (N2) were cultured using standard laboratory conditions on *E. coli* OP50 or HT115 expressing appropriate RNAi. RNAi expression was induced using 6 mM IPTG. Adults were bleached onto RNAi plates and allowed to develop to the L4 to young adult transition before stresses were applied. For heat stress applications, animals were raised at 15°C from hatching then at the L4/young adult transition replicate plates were placed at 15°C or 37°C for 2 hours. After each stress, animals were washed off the plates with S-basal, then pellets frozen at −80°C. RNA was prepared as in^12^. For survival assays, ~10 −15 adult N2 animals were bleached on 60 mm RNAi plates. The eggs were allowed to hatch and grow to young adults at 15°C. 25-30 young adults were then moved to 35 mm plates in triplicate (75-90 animals per RNAi treatment) and subjected to heat shock at 37°C for 2 hours. Animals were kept at 20°C for the remainder of the assay. Dead animals were identified by gentle prodding, were counted and removed each day. Animals that died of bagging or from desiccation on the side of the plate were not counted. Three independent non blinded biological replicates were carried out and Kaplan-Meir curves were generated with GraphPad Prism v8.0. For lifespan experiments, the N2 adults were bleached on 60 mm RNAi plates. The eggs were allowed to hatch and grow to young adults at 20°C. 25-30 young adults were then moved to 35 mm plates in triplicate (75-90 animals per RNAi treatment). Adults were moved to fresh plates every day and dead animals were identified by gentle prodding and removed each day. Three independent non blinded biological replicates were carried out and Kaplan-Meir curves were generated with GraphPad Prism v8.0.

### Gene expression analysis, RNA sequencing and analysis

RNA for deep sequencing was purified by Qiagen RNAeasy. Duplicate samples were sent for library construction and sequencing at BGI (China). Raw sequencing reads were processed using an in-house RNA-Seq data processing software Dolphin at University of Massachusetts Medical School ^63^. The raw read pairs were first aligned to *C. elegans* reference genome with ws245 annotation. The RSEM method was used to quantify the expression levels of genes and Deseq was used to produce differentially expressed gene sets with more than a 2-fold difference in gene expression, with replicates being within 0.05 in a Students T test and a False Discovery Rate (FDR) under 0.01. Statistics were calculated with DeBrowser ^64^. Venn Diagrams were constructed by BioVenn ^65^. WormCat analysis was performed using the website www.wormcat.com^27,66^ and the whole genome annotation version 2 (v2) and indicated gene sets. PCA was conducted by using *prcomp* in R and graphed with *ggplot* in R studio.

### Immunofluorescence

For H3K4me3 (Cell Signaling Technology, catalogue number C42D8) staining, dissected intestines were incubated in 2% paraformaldehyde, freeze cracked, then treated with −20°C ethanol before washing in PBS, 1% Tween-20, and 0.1% BSA. Images were taken on a Leica SPE II at identical gain settings within experimental sets. Quantitation was derived for pixel intensity over nuclear area for at least seven dissected intestines, with at least 3 nuclei per intestine. Three biological repeats were carried out for every experiment.

### Sample preparation for LC-MS

*C. elegans* (N2) gravid adults (~15-20) were bleached onto 60 mm RNAi plates, eggs were allowed to hatch and grow to young adults at 15°C. For heat stress application, replicate plates were placed at either 15°C or 37°C for 2 hours. At the end of the heat stress, worms were collected in S-Basal, and pellets were frozen at −80°C. Four independent biological replicates were collected. To prepare the samples for LC-MS, the pellet was thawed on ice and washed with 0.9% NaCl. Washed worms were then transferred to 2 mL FASTPREP tubes (MP Biomedicals) containing 1.4 mm ceramic beads (Qiagen). The samples were then resuspended in 1 mL 80% methanol (LC-MS grade) and homogenized using a bead beater (6.5 m/s; 20 seconds). The samples were cooled on ice between cycles. The homogenized samples were then vortexed at 4°C for 10 min and centrifuged at 21,000 RPM for 10 min at 4°C. The supernatant was removed at dried under vacuum. The pellet was resuspended in ice cold RIPA buffer and vortexed at 4°C for 10 min and centrifuged at 21,000 RPM at 4°C for 10 min. The supernatant was removed and used for protein quantification using Pierce Protein BCA assay kit (ThermoFisher). The protein quantification was then used to resuspend the pellet for an equal input of 0.5 μg/ml of protein per sample.

### LC-MS analysis

#### Absolute quantification of SAM

Samples were extracted in 80% methanol containing 500 nM methionine-^13^C_5_-^15^N (Cambridge Isotope Laboratories, Inc.) as an internal standard and metabolites were detected as described above. Absolute quantification of SAM was performed using an external calibration curve prepared with synthetic standard, and peak areas were normalized to methionine-^13^C_5_-^15^N. Normalized peak areas from the standard curve were fit to a quadratic log-log equation with an r^2^ value of >0.995 which was then used to calculate the concentration of SAM in each sample. Statistical analysis was carried out for the data using GraphPad Prism (v8.0).

#### Relative metabolite profiling

Metabolomics was conducted on a QExactive Plus bench top orbitrap mass spectrometer equipped with an Ion Max source and a HESI II probe, which was coupled to a Vanquish Horizon HPLC system (Thermo Fisher Scientific, San Jose, CA). External mass calibration was performed using the standard calibration mixture every 7 days. Dried extracts were reconstituted in enough water to achieve a final concentration of 0.5 μg/ml protein per sample. 2 μL of this resuspended sample were injected onto a SeQuant^®^ ZIC^®^-pHILIC 150 × 2.1 mm analytical column equipped with a 2.1 × 20 mm guard column (both 5 mm particle size; Millipore Sigma). Buffer A was 20 mM ammonium carbonate, 0.1% ammonium hydroxide; Buffer B was acetonitrile. The autosampler tray was held at 4°C. The chromatographic gradient was run at a flow rate of 0.150 mL/min as follows: 0-20 min: linear gradient from 80-20% B; 20-20.5 min: linear gradient form 20-80% B; 20.5-28 min: hold at 80% B. The mass spectrometer was operated in full-scan, polarity-switching mode, with the spray voltage set to 4.0 kV, the heated capillary held at 320°C, and the HESI probe held at 350°C. The sheath gas flow was set to 10 units, the auxiliary gas flow was set to 1 units, and the sweep gas flow was set to 1 unit. MS data acquisition was performed in a range of *m/z* = 70–1000, with the resolution set at 70,000, the AGC target at 1×10^6^, and the maximum injection time at 20 msec. An additional scan (*m/z* 220-700) in negative mode only was included to enhance detection of nucleotides. Relative quantitation of polar metabolites was performed TraceFinder 5.1 (Thermo Fisher Scientific) using a 5 ppm mass tolerance and referencing an in-house library of chemical standards. Statistical analysis was carried out for the data using GraphPad Prism (v8.0).

### CUT&Tag

*C. elegans* (N2) were cultured using standard laboratory conditions on *E. coli* OP50. Adults were bleached onto RNAi plates and allowed to develop to the L4 to young adult transition before heat stress was applied. For heat stress applications, animals were raised at 15°C from hatching then at the L4/young adult transition replicate plates were placed at 15°C or 37°C for 2 hours. At the end of the heat stress, animals were washed off the plates with S-basal, then pellets frozen at −80°C. Worm pellets were washed with S-Basal to remove bacteria, then resuspended in 750 uL of chilled Nuclei Purification Buffer (50 mM HEPES pH =7.5, 40 mM NaCl, 90 mM KCl, 2 mM EDTA, 0.5 mM EGTA, 0.2 mM DTT, 0.5 mM PMSF, 0.5 mM spermidine, 0.1% tween 20, and cOmplete proteinase inhibitor cocktail (Roche)). The suspension was then transferred to Potter-Elvehjem Tissue Grinder (3 mL). The worms were ground with 2-3 cycles consisting of ~45-50 strokes of the grinder. The samples were chilled on ice for ~5 minutes between consecutive cycles. The lysates were passed through 100 micron filter (X3) followed by 40 micron (X3) (Pluriselect). The lysates were then centrifuged at 4500 RPM for 10 minutes at 4°C. The pellets were resuspended gently in wash buffer (1M HEPES pH 7.5, 5 M NaCl, 2 M spermidine). Concanavalin bead slurry (10 uL/sample) was added gently to the samples and allowed to incubate at room temperature for 15 min in an end-over-end rotator. The sample tubes were then transferred to a magnetic stand and liquid was gently removed. The nuclei were gently resuspended in 50 uL of chilled antibody buffer (8 μL 0.5 M EDTA, 6.7 μL 30% BSA in 2 mL Dig-wash buffer (400 μL 5% digitonin with 40 mL Wash buffer)). 1 uL anti-H3K4me3 antibody (Cell Signaling Technology, catalogue number C42D8) was added to the suspension and allowed to bind overnight at 4°C on a nutator shaker. Samples without any antibody added were used as controls to correct for background reads and further processed per the CUT&Tag protocol ^67^ to generate sequencing libraries. The libraries were amplified by mixing 21 μL of DNA with 2μL each of (10 μM) barcoded i5 and i7 primers, using a different combination for each sample. 25 μL NEBNext HiFi 2 × PCR Master mix (NEB) was added, and PCR was performed using the following cycling conditions: 72 °C for 5 minutes (gap filling); 98 °C for 30 seconds; 17 cycles of 98 °C for 10 seconds and 63 °C for 30 s; final extension at 72 °C for 1 minute and hold at 4 °C. 1.1 × volume of Ampure XP beads (Beckman Coulter) was incubated with libraries for 10 minutes at room temperature to clean up the PCR mix. Bead bound DNA was purified by washing twice with 80% ethanol and eluting in 20 μL 10 mM Tris pH 8.0. Size distribution of the libraries was determined by Fragment analyzer and concentration by the KAPA Library Quantification Kit before sequencing to determine the H3K4me3 landscape in basal and heat stress condition in worms fed on control, *sams-1* or *sams-4* RNAi. Sequencing of the prepared libraries was carried out on Illumina NextSeq 500.

### Data analysis

Paired end reads from each sample were aligned to the *C. elegans* genome (ce11 with ws245 annotations) using Bowtie2 ^68^with the parameters −N 1 and −X 2000. Duplicate reads were removed using Picard (http://broadinstitute.github.io/picard/) and the reads with low quality scores (MAPQ < 10) were removed. HOMER software suite was used to process the remaining mapped reads ^36^. The “makeUCSCfile” command was used for generating genome browser tracks. Data was normalized to library size. the “findPeaks <tag directory> -style histone -o auto” command was used for calling H3K4me3 peaks and the “annotatePeaks” command was used for making aggregation plots. Differential peak calling was accomplished using ^37^the command “. We used the findOverlapsOfPeaks command in ChipSeqAnno^37^ with a max gap of 1000 basepairs to determine peak overlap. TSS plots were generated using HOMER ^36^ and Venn Diagrams were constructed by BioVenn ^65^.

Correlation matrices were generated with deeptools version 3.5.1^69^. Multibamsummary was used to compare bam files from each sample, using default values except --binSize 2000. This data was visualized using plotCorrelation with --removeOutliers and the Pearson method. Previously published datasets were used to compare H3K4me3 Cut and Tag versus previously published data sets. Young adults fed a normal diet were used from Wan et al. 2022^40^. Day 2 *glp-1* adults were chosen from Pu et al^39^. modENCODE ChIP-seq data drew from L3 animals^38^.

## Acknowledgements

We would like to acknowledge the Walker lab for reading of the manuscript, Drs. Marian Walhout and Craig Peterson for helpful discussions and Life Science editors for manuscript assistance. Absolute quantification of SAM was carried out at the Whitehead Metabolomics Core (Cambridge, MA). We thank the UMASS Transgenic animal core (Dr. Paola Perrat and Dr. Michael Francis) for construction of RFP::SAMS-1and GFP::SAMS-4. Funding is from NIH: 1R01AG053355 to AKW and R01HD072122 to TGF.

## Supplemental data

**Fig S1: Expression patterns of SAM synthases in adult *C. elegans***

**(A)** Comparison of polyA+ RNA levels of SAM synthases with selected other metabolic genes in adult animals from the modEncode data set^24^. **(B)** Representative confocal images of animals expressing RFP::SAMS-1or GFP::SAMS-4. hypodermis is (h), intestine (i) and germline (gl). (**C**) Confocal projections of GFP::SAMS-4 and SAMS-3::mKate subjected to *sams-3* or *sams-4(RNAi)*. (**D**) Absolute quantification of the SAM level in animals fed on control RNAi or *sams-4(RNAi)*. The levels are expressed as mM/mg tissue.

**Figure S2: Distinct patterns of gene expression after *sams-1 or sams-4* RNAi in basal conditions**

**(A)** Principal component analysis showing overlapping components between genes regulated in *sams-3* and *sams-4(RNAi)* animals. **(B)** Venn diagram showing the overlap in up regulated genes in animals fed *sams-1* or *sams-4(RNAi)*. **(C)** Bubble charts show broad category enrichment of up regulated genes in animals fed *sams-1* or *sams-4(RNAi)*. **(D)** Bubble charts show broad category enrichment of down regulated genes in animals fed *sams-1* or *sams-4(RNAi)*. **(E)** Venn diagram showing the overlap in up regulated genes involved in lipid metabolism in animals fed *sams-1* or *sams-4(RNAi)*. **(F)** Venn diagram showing the overlap in up regulated genes involved in pathogen stress response in animals fed *sams-1* or *sams-4(RNAi)*.

**Fig S3: *sams-4* is important for survival and H3K4me3 in *sams-1* animals after heat shock.**

**(A)** Schematic for the heat stress assay. (**B**) Survival assays comparing response to heat in SAM synthase mutants. (**C**) Lifespan assay with *sams-4(RNAi)* animals where *sams-3* may also be targeted. (**D**) Heat shock survival assays showing that genetic loss of *sams-4* limits survival in *sams-1(RNAi)* animals after heat shock. For B-D, statistical significance is shown by Log-rank test. Each graph represents compiled data from 3 biologically independent repeats. Data for each replicate is compiled in Table S2. **(E)** Quantification of immunofluorescence imaging of intestinal nuclei stained with HK4me3 antibody after heat shock from *sams-4(ok3315); sams-1(RNAi)* animals. Statistical significance was calculated using unpaired Student’s t-test. ns= not significant, **** = p<0.0001, *** = p<0.001. Graph represents compiled data from three biologically independent repeats per condition. LC/MS relative quantitation of SAM (**F**), Methionine (**G**) and S-adenosylhomocysteine (SAH) (**H**). Graphs represent 4 independent biological replicates (1-4: red, blue, orange and green) that were normalized for protein levels before quantitating relative levels of metabolites.

**Figure S4: H3K4me3 demethylases modulate SAM synthase phenotypes**

Representative immunofluorescence images and quantitation of intestinal nuclei stained with H3K4me3 specific antibody for *set-2* (**A, D**), *set-16* (**B, E**) and *rbr-2* (**C, F**). Statistical significance was calculated using unpaired Student’s t-test. ns= not significant, **** = p<0.0001, *** = p<0.001. Graph represents compiled data from three biologically independent repeats per condition. Heat shock survival assays examining the impact of demethylase knockdown on *sams-1(lof)* animals for *amx-1* (**G**) and *lsd-1* (**H**). Survival was determined by plotting Kaplan-Meier survival plots. Statistical significance is shown by Log-rank test. Each assay represents compiled data from 3 biologically independent repeats (**Table S2**).

**Figure S5: H3K4me3 C&T correlation with published H3K4me3 ChipSeq data.**

(**A**) Correlation plots showing r values for C&T replicates. (**B**) Comparison of H3K4me3ChIP seq from modEncode (L3)^38^, Pu et al (Adult *glp-1(e2141))^41^*, Wan et al (adult)^40^ and our C&T data. (C) IGV browser tracks showing no antibody controls around the pcaf-1 gene, which has been used as positive control for H3K4me3 5 prime peaks in *C. elegans^12,32^*.

**Figure S6. Distinct gene expression and H3K4me3 patterns after heat shock in *sams-1* and *sams-4* RNAi animals**

Sunburst diagram showing the enriched gene categories in animals fed control RNAi at **(A)** 15°C or **(B)** 37°C. Sunburst diagram showing the overall enriched gene categories **(C)** and genes involved in metabolism **(D)** in animals fed *sams-1(RNAi)* at 37°C. Sunburst diagram showing the overall enriched gene categories **(E)** and genes involved in metabolism **(F)** in animals fed *sams-4(RNAi)* at 37°C. Aggregation plots showing average enrichment of reads around the transcription start site (TSS) for genes which are *sams-1* dependent only dependent on either *sams-1* or *sams-4* or *sams-4* dependent only at **(G)** 15°C or **(H)**37°C. The Y axis on TSS plots shows Peaks per base pair of gene.

**Fig S7. SAM synthase-specific patterns H3K4me3 in germline nuclei.**

**(A)** Representative immunofluorescence images of H3K4me3 staining in the germline in animals fed on control, *sams-1* or *sams-4(RNAi)*. **(B)** Venn diagrams showing the overlap in H3K4me3 peaks identified on ubiquitously expressed genes in control animals at 15°C or 37°C. **(C)** Venn diagrams showing the overlap in H3K4me3 peaks identified on germline specific genes in control animals at 15°C or 37°C. Aggregation plots showing average enrichment of reads around the transcription start site (TSS) of **(D)** ubiquitously or **(E)** germline specific or **(F)** intestine specific genes in animals fed control, *sams-1* or *sams-4(RNAi)* at 15°C. The Y axis on TSS plots shows Peaks per base pair of gene. Aggregation plots showing average enrichment of reads around the transcription start site (TSS) of **(G)** ubiquitously or **(H)** germline specific or **(I)** intestine specific genes in animals fed control, *sams-1* or *sams-4(RNAi)* at 37°C.

**Figure S8: *sams-1* and *sams-4* have distinct gene expression patterns after heat shock.**

(**A**) PCA plot showing groupings of up and down regulated genes from Control, *sams-1* or *sams-4(RNAi)* animals. Survival curves examining heat shock responses after RNAi of T27F6.8 or *fbxa-59*. Survival was determined by plotting Kaplan-Meier survival plots. Statistical significance is shown by Log-rank test. Each assay represents compiled data from 3 biologically independent repeats (**Table S2**).

**Figure S9: Schematic of potential *nhr-68* module regulation in sams-1 animals.**

(**A**) Genome browser tracks for *ges-1* showing H3K4me3 enrichment in animals fed control, *sams-1* or *sams-4(RNAi)* at 15°C or 37°C. (**B**) Schematic showing the dynamic changes in the transcription and H3K4me3 landscape in low SAM animals following heat shock.

**Table S1 (Microsoft Excel File): RNA seq for SAM synthase knockdown in basal conditions. Tabs A-C** show *sams-3, sams-4, sams-5 (RNAi)* RNA seq data then **Tabs D-F** show WormCat gene enrichment. *sams-1* data is from Ding, et al. 2018. Enriched categories from WormCat. Red color denoted categories with a p value of less than 0.01. NS is not significant, NV is no value, RGS is regulated gene set.

**Table S2 (Microsoft Excel File) Statistics for survival curves.** Each tab contains data for replicate experiments (R1, R2, R3). Statistical information from GraphPad Prism is also included.

**Table S3 (Microsoft Excel File). Tabs A-F:** Cut and Tag peaks from Control, *sams-1* and *sams-4* RNAi animals at 15 and 37 degrees determined by HOMER. **Tabs G-I:** Enriched categories from WormCat. Color denoted categories with a p value of less than 0.01 NS is not significant, NV is no value, RGS is regulated gene set.

**Table S4 (Microsoft Excel File). Limited activation of heat shock response in *sams-4* RNAi animals.** Tabs show RNA seq from control (**A**), *sams-1* (**B**) *or sams-4* **(C**) animals subjected to heat shock that was used for comparison with C&T data. Differential genes were identified using Deseq2 in DolphinNext. Data for control and *sams-1* RNAi animals is from Ding, et al 2018. WormCat batch output of two-fold regulated genes for Categories 1, 2 and 3 are in tabs E-G). Highlighting denotes genes with significantly p values. NS is not significant, NV is no value, RGS is regulated gene set.

