## Supplementary figures and images for "S-adenosylmethionine synthases specify distinct H3K4me3 populations and gene expression patterns during heat stress"

### Figure S1

Godbole et al
Fig S1

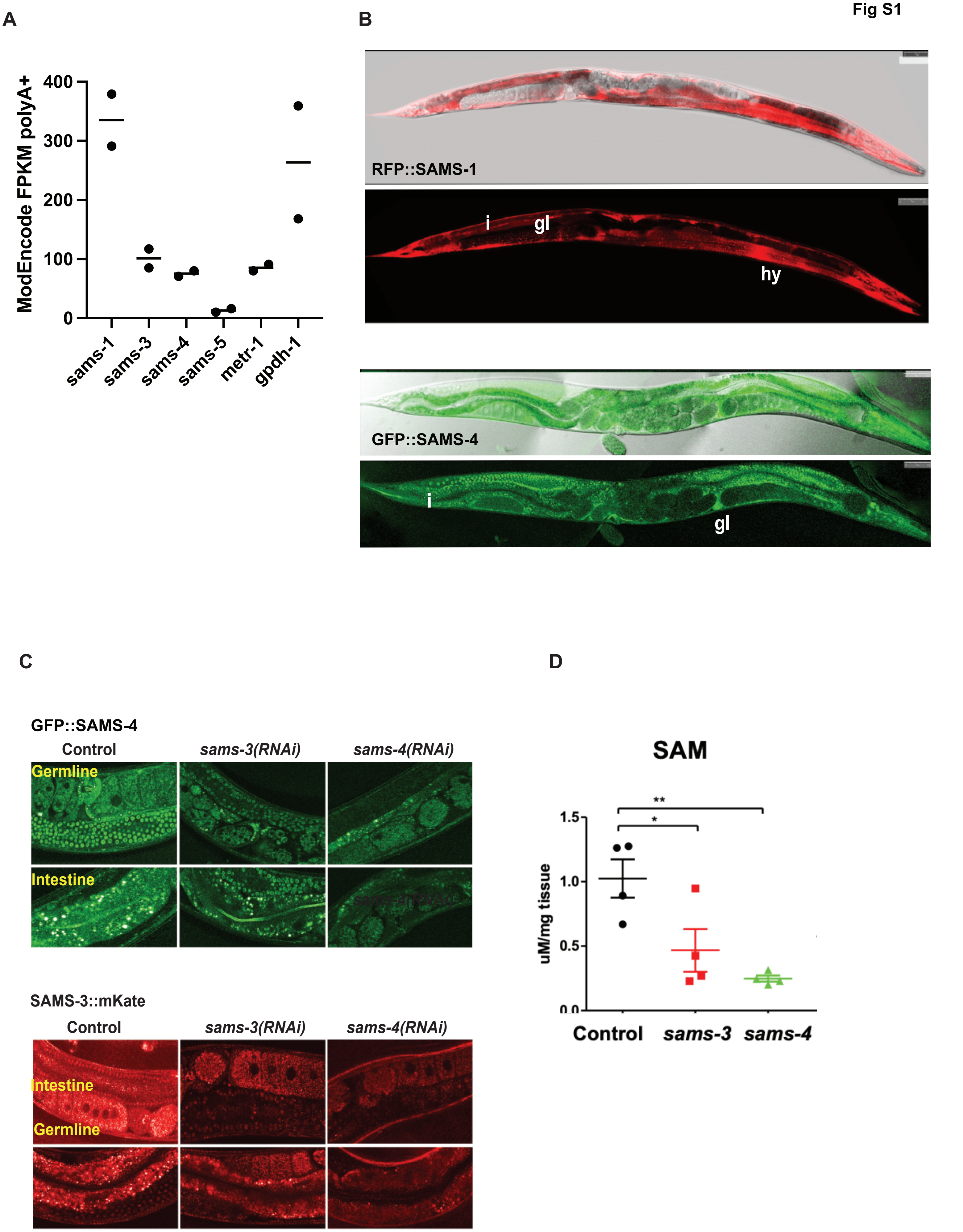

### Figure S2

Godbole, et al.
Fig S2

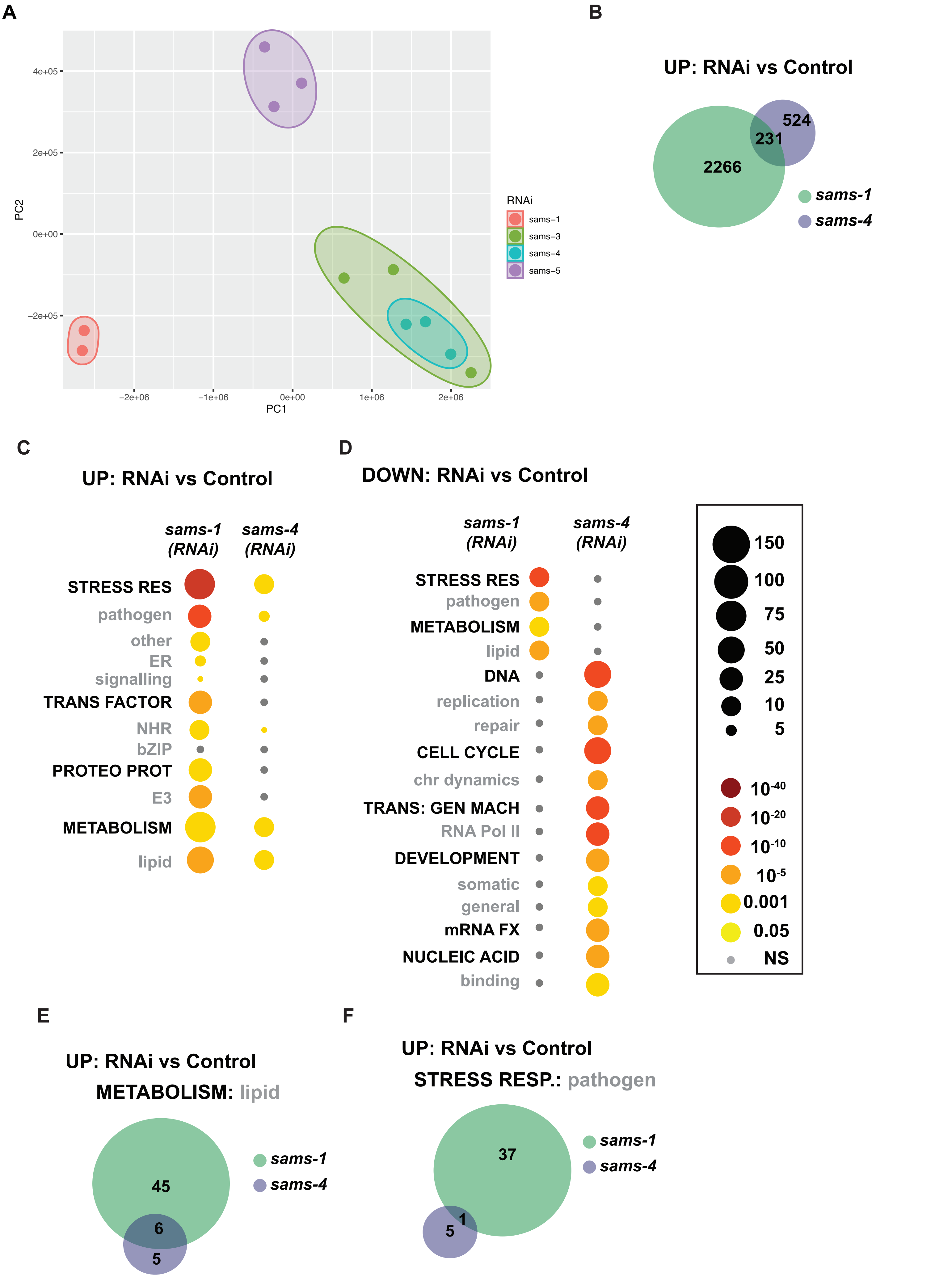

### Figure S3

Godbole, et al.
Fig S3

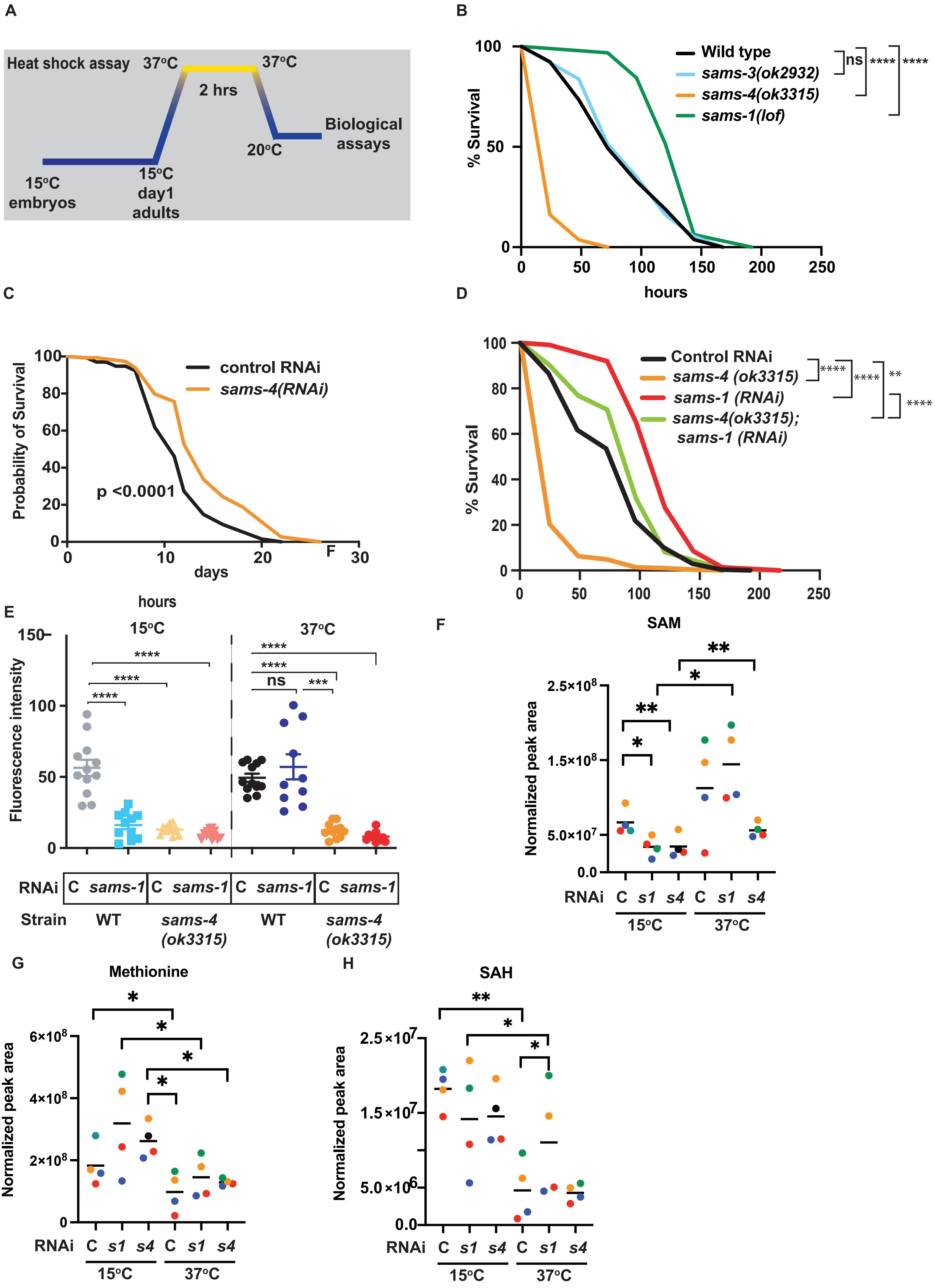

### Figure S4

Godbole et al
Fig S4

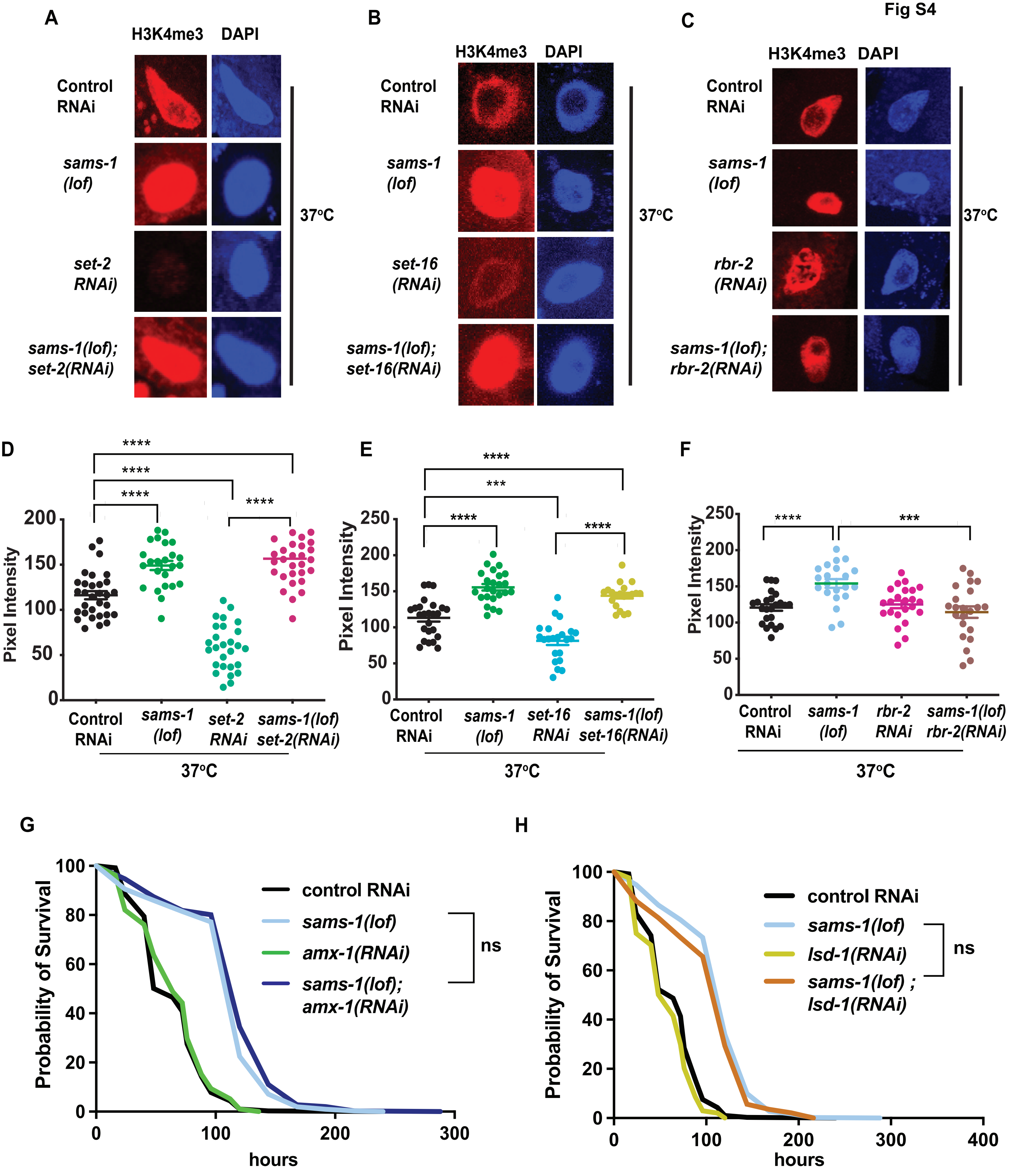

### Figure S5

Godbole et al
Fig S5

A

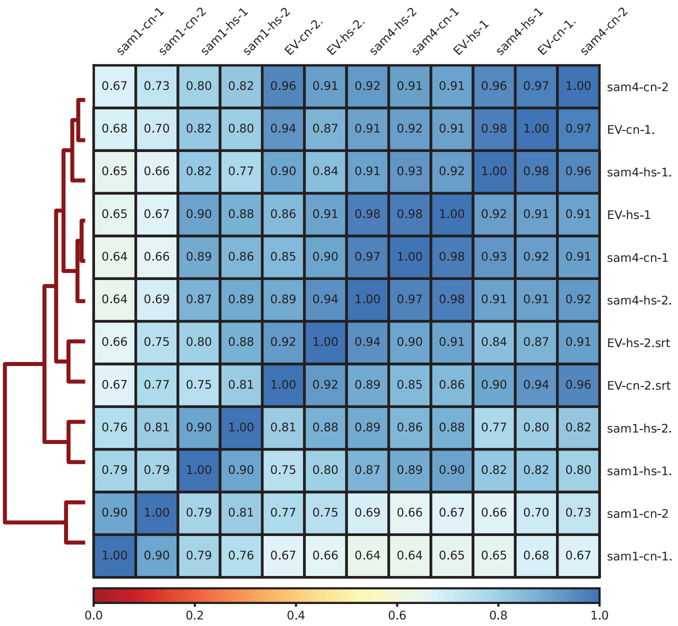


B

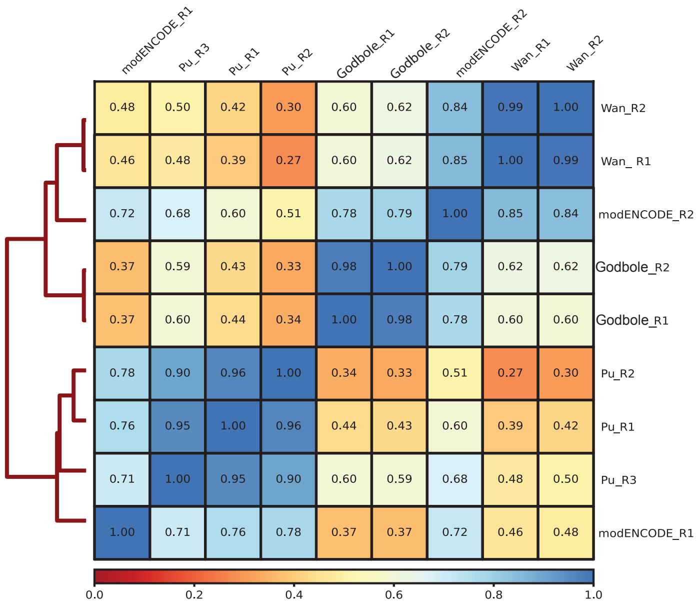


C

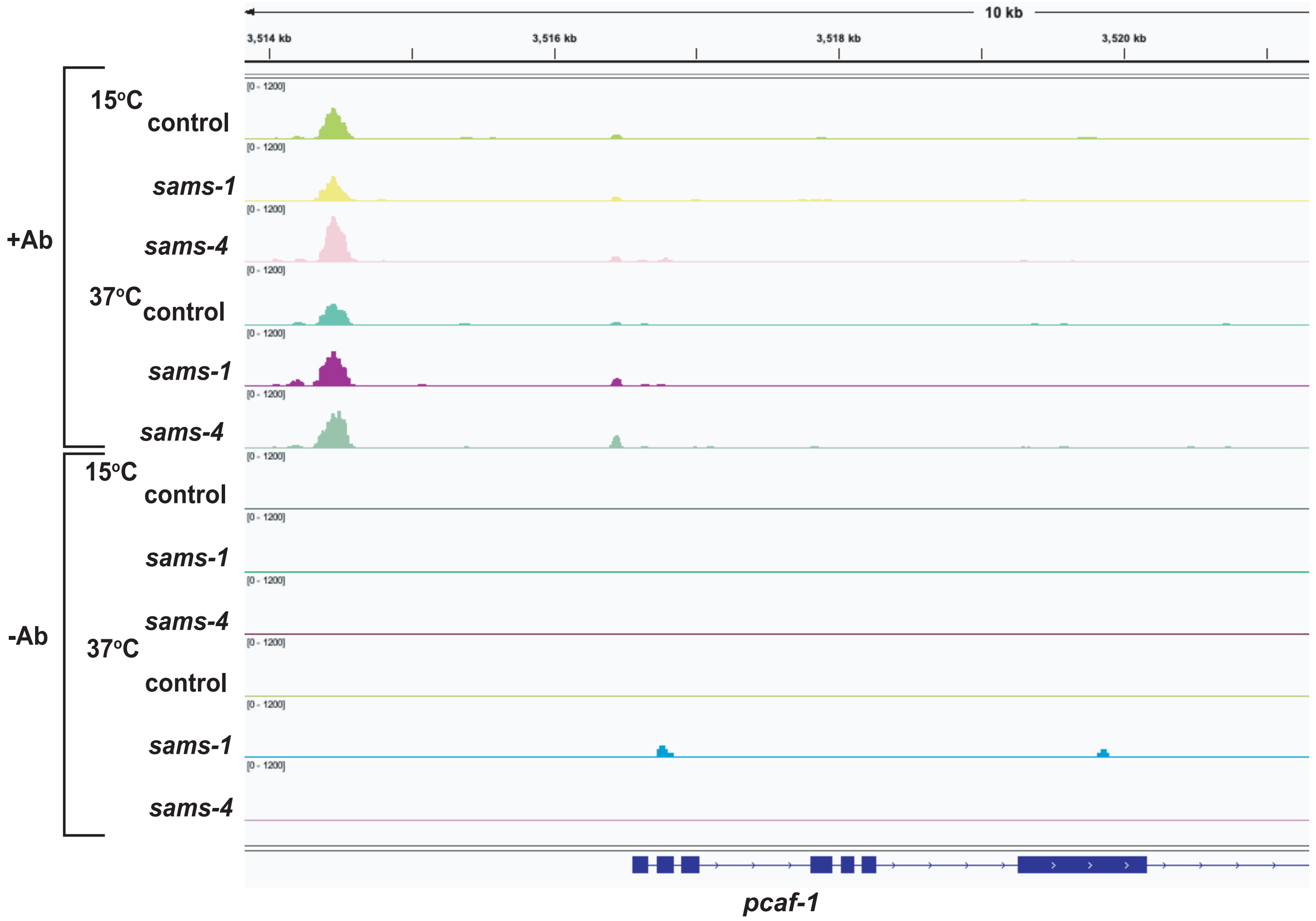

### Figure S6

Godbole et al
Fig S6

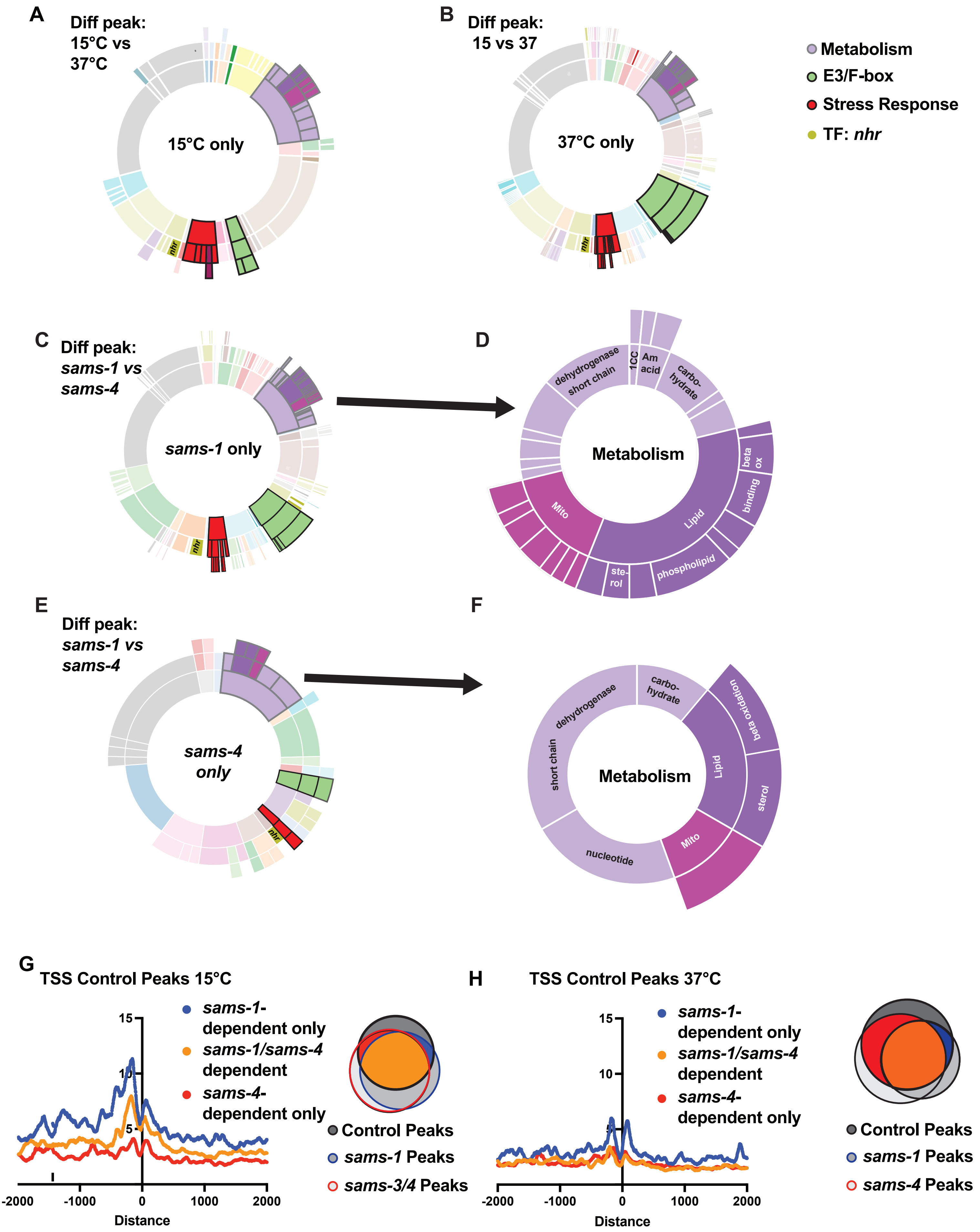

### Figure S7

Godbole et al
Fig S7

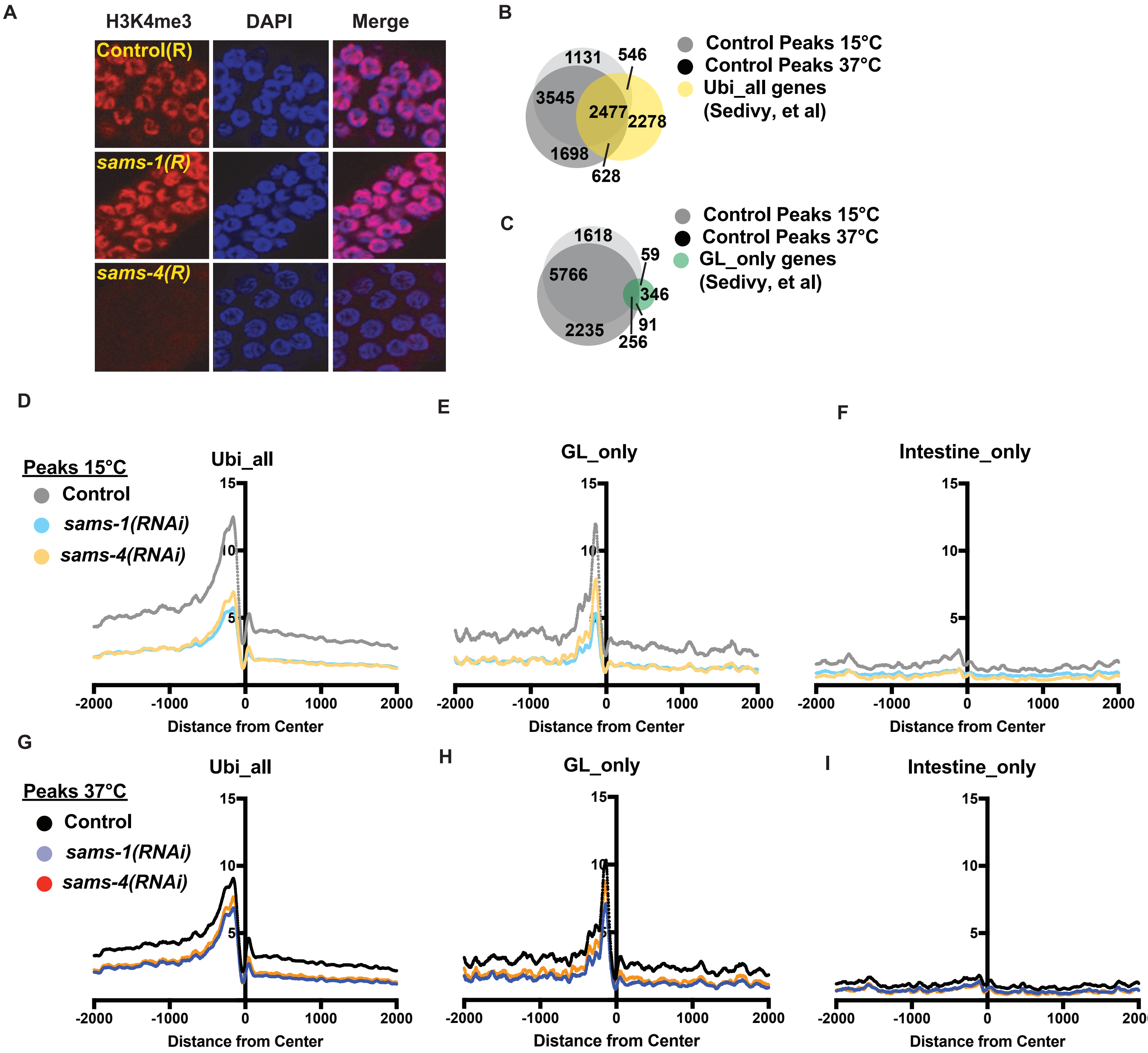

### Figure S8

**Godbole et al**
**Fig S8**

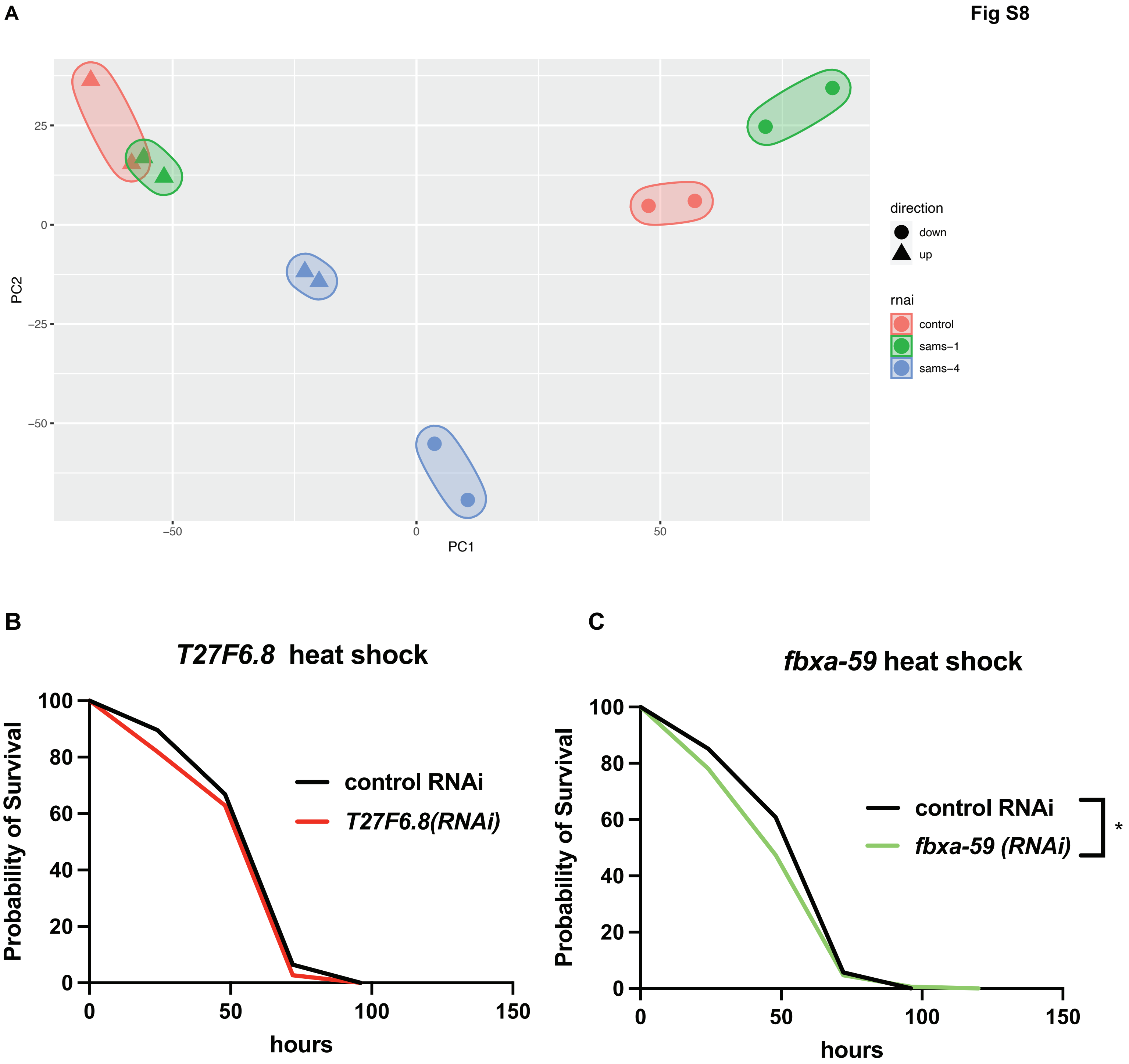

### Figure S9

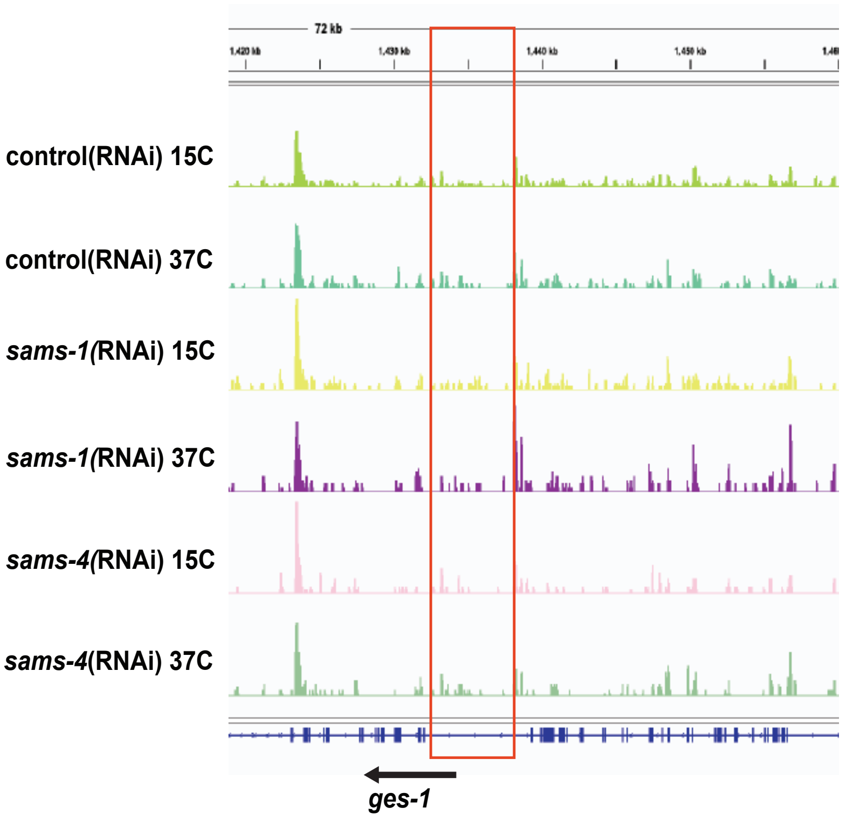

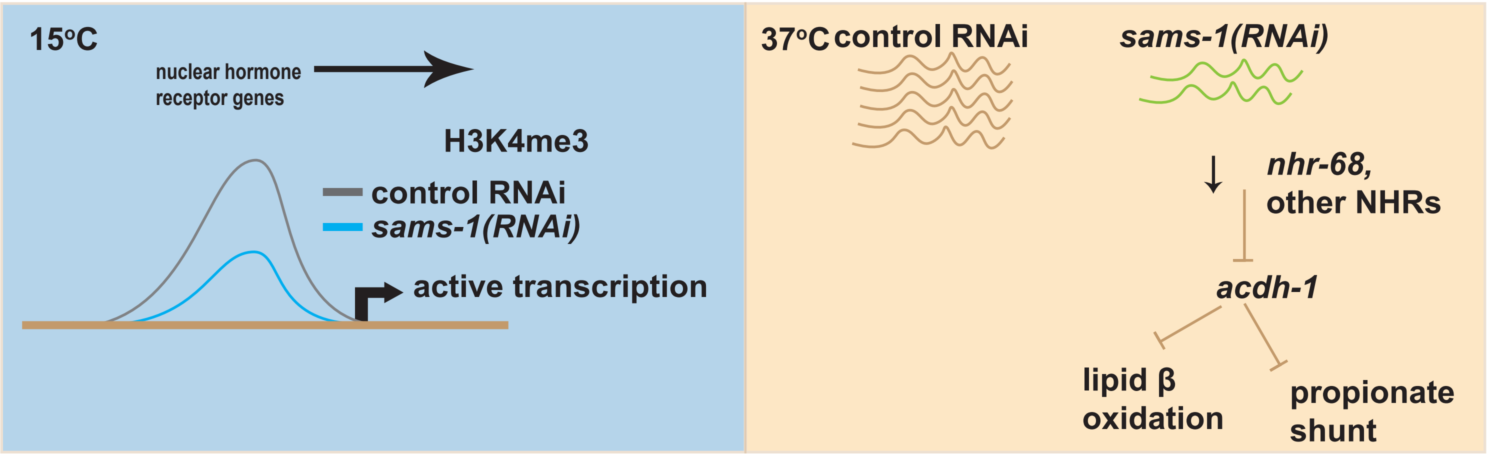
